# Viral chronotypes and their role in shaping seasonal viral dynamics in the Western English Channel

**DOI:** 10.1101/2024.05.16.594516

**Authors:** Luis M. Bolaños, Michelle Michelsen, Ben Temperton

## Abstract

Marine viruses are key players of ocean biogeochemistry, profoundly influencing microbial community ecology and evolution. Despite their importance, few studies have explored the temporal dynamics of viral genome abundances in marine environments. Viral dynamics are complex, influenced by multiple factors such as host population dynamics and environmental conditions. To disentangle the complexity of viral communities, we developed an unsupervised machine learning framework to classify viral genomes into “chronotypes” based on temporal abundance patterns. Analysing an inter-seasonal monthly time-series of surface viral metagenomes from the Western English Channel, we identified chronotypes and compared their functional and evolutionary profiles. Results revealed a consistent annual cycle with steep compositional changes from winter to summer and steadier transitions from summer to winter. Seasonal chronotypes were enriched in potential auxiliary metabolic genes like ferrochelatases and 2OG-Fe(II) oxygenases compared to non-seasonal types. Chronotypes clustered into four groups based on their correlation profiles with environmental parameters, primarily driven by temperature and nutrients. Viral genomes exhibited a rapid turnover of polymorphisms, akin to Red Queen dynamics. However, within seasonal chronotypes, some sequences exhibited annual polymorphism recurrence, which declined over a 16-month period, suggesting that a fraction of the seasonal viral populations evolve more slowly. Classification into chronotypes revealed viral genomic signatures linked to temporal patterns, likely reflecting metabolic adaptations to environmental fluctuations and host dynamics. This novel framework enables the identification of long-term trends in viral composition, environmental influences on genomic structure, and potential viral interactions.

## Introduction

Viruses are the most abundant entities in the ocean with a vast and mostly unexplored genetic functionality [1–3]. Plankton communities are the primary drivers of global biogeochemical cycles [4]. These cycles comprise primary producers and heterotrophic microorganisms that fuel oceanic trophic networks [5], the fluxes of biogenic compounds to and from the atmosphere [6] and their export to the deep sea [7,8]. Therefore, viruses that infect planktonic microorganisms in the ocean play important roles in global biogeochemical cycles by gatekeeping the flux of dissolved organic matter [9,10] or encoding auxiliary metabolic genes (AMGs) to modify hosts metabolism, impacting processes such as carbon, nitrogen, sulphur, and iron cycling [11–14].

The complexity of the potential outcomes and the environmental forces that control the viral-plankton interactions have been studied in a few systems. Examples include the viral enhancement of host fitness in Antarctic bacterial communities during low-nutrient seasons [15], the increase in phytoplankton mortality and viral production in the stratified summer North Atlantic [16], and the decrease in viral particles along the nutrient gradient towards oligotrophic conditions [17–20]. Despite these advances, a more detailed ecological and evolutionary understanding of the relationship between viruses, hosts and environmental conditions is needed to improve our modelling capacity and predict the warming impacts on oceanic ecosystems [21].

Metagenomics and metatranscriptomics emerged as a cost-effective methodology to analyse viral communities in large scales. Since the retrieval of molecular datasets from the field, it has become evident that viral diversity is immense [22]. Multiple field molecular surveys have expanded our knowledge on the biogeographic distribution of oceanic dsDNA phages [23,24], the abundance and distribution of RNA viruses [25], and the biogeographical patterns of macrodiversity and microdiversity [3]. Field metagenomes have revealed patterns in the genomic population boundaries and the ecological zones in which viral communities can be classified. However, phage-host relationships are shaped in a rapid evolutionary arms race [26], from which transect metagenomes can only offer snapshots of specific time points at different locations.

Oceanographic observatories provide a unique opportunity to overcome this limitation by generating viral metagenomic (virome) time-series. A handful of longitudinal viromes have been generated, providing important insights into seasonal succession, phage-host interactions, molecular evolution, and the overall environmental influence on viral composition [27–29]. In different locations and regimes, viral communities show a remarkably stable annual seasonality of populations (vOTUs) in both abundance [30] and composition [27,29,31]. Despite the annual stability of vOTUs, the genotypic configuration within populations undergoes constant turnover [27]. This resembles the dynamics outlined in the Red Queen Hypothesis, in which despite a continuous evolution, population fitness remains unchanged [32]. Recently, it has been shown that cyanophages display distinct temporal dynamics driven by the seasonality of the hosts, stochastic environmental processes, and different host range strategies [33]. However, these temporal patterns have yet to be explored in the context of the overall viral community to define how environmental pressures collectively influence them and whether potential interactions between them exist. Furthermore, an accurate classification of temporal abundance patterns of thousands of viral sequences within a time series remains a major methodological challenge. Generating these classifications represent a milestone in elucidating the associations between the distinct temporal abundance profiles of viral populations and specific gene functions, evolutionary patterns, and bottom-up environmental controls.

In this study, we generated a monthly virome times-series of the surface Western English Channel (WEC) to analyse the temporal dynamics of viral populations and the selection of viral genes influenced by seasonal environmental conditions. The WEC is a temperate coastal sea that stratifies seasonally during the summer and is highly dynamic due to the constant riverine input and wind mixing [34]. The WEC microbial community composition has been extensively studied [35–38], providing a perfect framework to analyse the virome dynamics. By developing a novel unsupervised machine-learning method, we categorized the temporal abundance patterns of genomes from vOTUs representative sequences into chronotypes. Chronotypes are statistically discrete clusters of co-varying genomes with similar temporal abundance patterns and optimally separated from other clusters within a dataset. This categorization allowed us to discern between viral populations with specific seasonal and non-seasonal patterns, interrogate whether viral genes are enriched in different temporal patterns, analyse the influence of environmental conditions on chronotypes and evaluate whether distinct evolutionary patterns exist. Our study provides the first detailed insights of viral dynamics in the WEC and their response to environmental variations. Furthermore, we present a novel method of temporal classification that could potentially be applied to analyse other relevant longitudinal molecular datasets, such as amplicon surveys or metatranscriptomes.

## Materials and Methods

### Sampling, fractionation, viral DNA extraction and sequencing

20 L of water samples were collected monthly from the surface (1 m) of the station L4 in the WEC (50°15N, 4°139W) (Table S1). Metadata was retrieved from the Western Channel Observatory [39] and the Archive for Seabed Species and Habitats (DASSH) [40]. Chlorophyll *a* concentrations were retrieved from the British Oceanographic Data Centre NOC [41]. Water samples were processed as described previously [42]. Briefly, viral enrichment was done by filtering sequentially the seawater through a glass fibre (GF/D: pore size 2.7 µm) and polyethersulfone (0.22 µm) filters in a 142 mm polycarbonate rig. The viral particles were precipitated using iron chloride and collected on a 1.0 µm polycarbonate filter [43]. Viruses were resuspended in ascorbate-EDTA buffer (0.1 M EDTA, 0.2 M MgCl_2_, 0.2 M ascorbic acid, pH 6.0) using 2 mL of buffer per 1 L of seawater. The resuspended viral fraction was transferred to an Amicon Ultra 100 kDa filter, pre-treated with 1% bovine serum albumin and flushed with SM buffer (0.1 M NaCl; 0.05 M Tris–HCl; 0.0008 M MgCl_2_) [44]. The viral fraction was concentrated to 600 µl and dissolved DNA was digested using DNAse1 (100 U/mL; 2 h at room temperature). DNA was extracted using the Wizard DNA Clean-Up System (Promega A7280) and cleaned with a 1.5× Ampure bead cleanup. 50 ng of DNA were used to construct sequencing libraries in a 1S Plus (Swift Biosciences) library preparation. Libraries were sequenced on an Illumina HiSeq 2500 (2×125 bp) to a depth of ∼25M at the University of Exeter sequencing facility.

### Viromes processing, assembly, identification, gene annotation, and vOTUs definition

All datasets were processed individually following the protocol described in [3]. Briefly, quality processing was done with BBtools v38.96 [45]. Sequencing adaptors and ΦX174 reads were removed using bbduk (ktrim=r,k=23,mink=11,hdist=1,minlen=50). Reads that mapped to the human genome were removed using bbmap (minid=0.95,maxindel=3,bwr=0.16,bw=12,quickmatch,fast,minhits=2). Short reads were assembled using metaSPAdes from SPAdes v3.15.5 [46]. Contigs with a minimum length of 10,000 bp were used as input to VirSorter v2.2.3 [47]. Contigs classified as viral were concatenated and clustered into viral operational taxonomic units (vOTU; >95% nucleotide identity and > 85% coverage) using the Perl script Cluster_genomes_5.1.pl (https://github.com/simroux/ClusterGenomes). The quality of the vOTUs was assessed with CheckV [48]. A final dataset of 3,090 high- and medium-quality vOTUs were clustered along with viral genomes from the INPHARED [49] database (July 2023) using vConTACT2 v0.11.1 [50]. Representative genomes of vOTUs were annotated using Pharokka v1.3 [51] and DramV v1.3.5 [52].

### Estimation of vOTUs relative abundance

To estimate the relative abundance of vOTUs, quality filtered virome reads were mapped to the representative genomes with bowtie2 v2.3 with default parameters [53]. The generated read alignment files were used to estimate the Reads Per Kilobase per Million mapped reads (RPKM) measurement for each vOTU representative genome of all samples using coverM v0.6.1 [54] with the following parameters --min-read-percent-identity 0.9 --min-covered-fraction 0.4. A Snakemake (v 5.26.0; [55]) file used to process the viral metagenomes, metadata, and intermediate processing products are available at https://github.com/lbolanos32/WEC_Chronotypes_2024.

### Unsupervised machine learning framework to generate chronotypes

Seasonality, defined as the annual periodic fluctuations of each vOTU, was determined with a Fisher G-test [56] implemented in the GeneCycle package v1.1.5 [57] and based on the RPKMs of the representative genomes. Seasonal and non-seasonal RPKM profiles were analysed independently using a developed clustering method: ChronoClustR (Fig. 1). Briefly, we implemented an iterative unsupervised clustering strategy in which z-score normalized values of the RPKM longitudinal profiles were randomly divided into subgroups of 100 elements. Each of these subgroups was clustered with TSclust v1.3 [58] to calculate the centroids of the optimal number of clusters of each subgroup (defined by having the best Silhouette score) based on the Euclidean distances between them. These centroids were then clustered using the K-means method for all potential numbers of clusters (from 1 to n-1 clusters, n=total number of centroids). Based on the curve generated of the relationship between the total within-clusters sum of squares and the number of clusters (*k*), we defined the optimal number of clusters by estimating the closest value of k to the first derivative (slope of the tangent line). Three types of clustering (kmeans, TSclust, and hierarchical clustering) were generated using the optimal *k* on the original collection. TSclust was selected for this study. These steps were bootstrapped to generate a distribution of the co-occurrence frequency of the vOTU longitudinal profiles in a cluster after multiple rounds of unsupervised clustering. In the current study, we performed 100 bootstrap resamples for both collections seasonal vOTUs and 100 for non-seasonal. A dendrogram was created based on the pairwise co-occurrence matrix of all vOTUs. Finally, the dendrogram was divided using the highest optimal *k* value estimated from all the iterations (see Supplementary methods). The method assumes equidistant sampling time-points with no missing values. Therefore, datapoints were labelled to the closest equidistant time-point within the series and missing values were interpolated using zoo package v1.8 [59]. R scripts and data used to generate chronotypes are available at https://github.com/lbolanos32/WEC_Chronotypes_2024.

**Figure 1.**
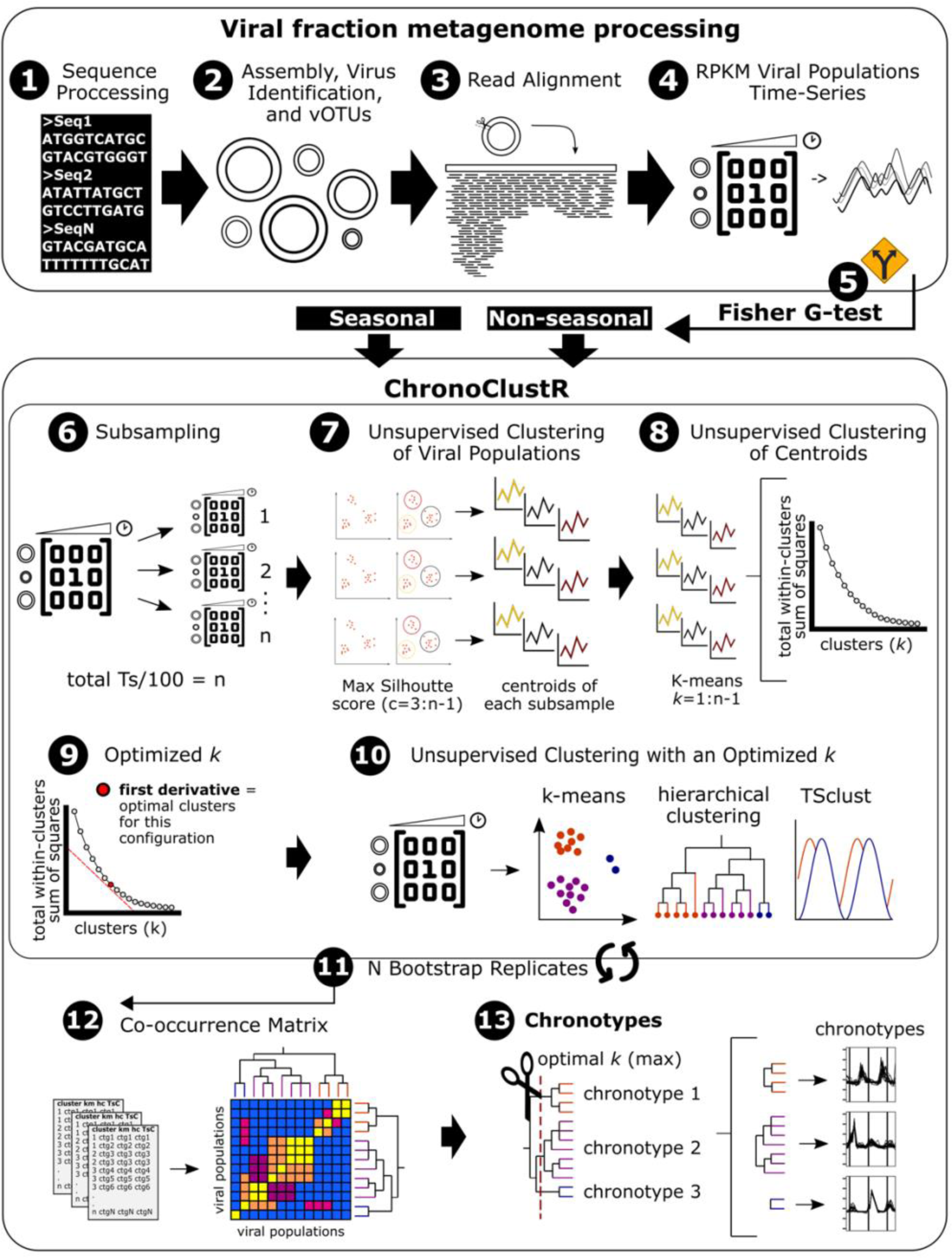
ChronoClustR: a workflow to generate highly defined time-series chronotypes using an unsupervised machine learning algorithm. **(1-4)** Generate a vOTU RPKM matrix organised as a function of time: raw metagenomic datasets are quality filtered and decontaminated, assembled, classified as viral, dereplicated into vOTUs, and quality filtered. RPKMs were generated from the short-read alignment to the quality-filtered vOTUs. **(5)** Determine the seasonal and non-seasonal vOTU time-series. **(6-10)** Single round of unsupervised clustering for seasonal and non-seasonal vOTU time-series. The inputted collection of vOTUs time-series is subsetted into groups of 100 time-series (default) without replacement, each of these groups is clustered based on the Euclidean distance between them using the R package TSclust (selecting the *k* that outputs a maximum Silhouette score; being the *k*’s tested 3 to maximum number of elements −1), the generated centroids of all subgroups are subjected to a new clustering round using k-means for each of all the potential number of *k* clusters (minimum = 1 and maximum = total centroids −1). Based on the relationship between the number of clusters and the total within-clusters sum of squares, the program estimates an optimal *k* using the first derivative. The original dataset is then reanalysed setting the optimal *k* generated in the previous steps. The workflow can do this clustering step using k-means, hierarchical clustering or TSclust. **(11-13)** Bootstrapping and chronotype definition. Steps 6 to 10 are bootstrapped (default = 100 replicates) to generate a matrix of co-occurence frequencies. This co-occurence matrix is used to generate a dendrogram which is then splitted into the maximum optimal *k* obtained through all bootstrap replicates (default). Each of the resulting groups will be considered a “chronotype”.

### Bray-Curtis dissimilarity and environmental Euclidean distances

Averaged Bray–Curtis dissimilarities were used to generate time-decay curves [60,61]. Euclidean distances between samples were generated based on normalized environmental measurements, and wavelet transforms of both measurements were done on WaveletComp [62] following previously described methods [63]. Bray-Curtis dissimilarity time-decay curves were generated for both seasonal and non-seasonal subsets. To identify whether the seasonal signal was driven by the most abundant vOTUs, we further divided seasonal and non-seasonal fractions into the first quartile, interquartile range, and third quartile of the RPKM distribution. A harmonic linear sinusoidal regression was generated in R v4.3 [64] and evaluated to confirm a significant seasonal amplitude in the data.

### Differential gene abundances and medoid correlations with environmental parameters

To assess the differential gene abundances between seasonal and non-seasonal chronotypes, we used the normalized values obtained by dividing the gene frequency by the number of members within a chronotype. Two-tailed Student’s t-tests with Bonferroni correction for multiple testing were performed in R v4.3 [64]. Spearman pairwise correlations were calculated for all combinations of chronotype medoids and environmental parameters. Corr.test function in Psych v2.3.9 [65] was used to report the probability values of each pairwise comparison (holm method for adjustment of multiple tests).

### Phylogenetic analysis

Predicted proteins belonging to significant differentially abundant gene categories between seasonal and non-seasonal chronotypes were analysed. To determine whether the enrichment of potential AMGs reflects a particular adaptation of a majority of seasonal pelagiphages (enriched in co-ocurring ferrochelatase and 2-OG(FeII) oxygenases) [66] or a broader observation, we analysed together both the vOTU proteins and those annotated from known pelagiphage genomes. Genomes from cultured and curated metagenome assembled pelagiphages [66–71] were re-annotated as previously described to standardise functional prediction. Predicted proteins annotated as ferrochelatase, primase, or 2-OG(FeII) oxygenases were retrieved and aligned together using t-coffee v12.0 [72]. Phylogenetic trees were constructed using IQ-Tree v2.2 [73] with the following parameters -msub viral -bb 1000 -mset. The best model selected for each category was: rtREV+F+R5 for ferrochelatase, rtREV+F+R7 for primase, and rtREV+F+I+R9 for 2-OG(FeII) oxygenases. Newick files were processed in R using ape v5.7 [74] and plotted with ggtree v3.8 [75].

### Polymorphism profiles

Single nucleotide polymorphisms were called as previously described [27]. Briefly, variation per site was estimated using samtools [76] and bcftools [77] on a subset of completely aligned reads with a minimum identity of 98% with the following parameters: ‘mpileup -g -f | bcftools call –ploidy 1 -mv’. Variants were filtered using ‘filter -e ‘%QUAL<20 || DP<10’; with a frequency >1%. The temporal variation profiles for 26,851 vOTUs longer than 10,000 bp were generated by counting shared polymorphisms between dates. No cross-assembly was performed to avoid potential chimeras and keep a consistent dataset of vOTUs. Two analyses were done, (a) considering only those vOTUs that had a minimum coverage of 10× across 90% of its length throughout all the samples [27] and (b) considering vOTUs that had a minimum coverage of 10× across 90% in at least one sample. Only 56 (out of the 3090 high-quality) vOTUs fulfilled these requirements in the analysis (a). These vOTUs represent a ubiquitous and abundant collection of viral sequences in the time-series. 2997 out of the 3090 high quality vOTUs fulfilled the conditions for analysis (b). Variant density was calculated for both sets as follows: 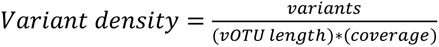. SNP profiles were compared using custom bash scripts. Figures were plotted using ggplot2 v2.3 package [78]. Figures were edited in Inkscape2 (www.inkscape.org) for aesthetics. R and bash scripts to generate the figures of this manuscript are available at https://github.com/lbolanos32/WEC_Chronotypes_2024.

## Results

### Western English Channel vOTUs composition is remarkably stable through time

A total of 26,851 vOTUs were generated from 27 monthly surface metagenomes spanning more than two and a half years in the WEC. Out of these, 3,090 (11.5%) were classified as high-quality viral genomes. Only nine (0.3%) of these high-quality vOTUs were classified as ssDNA phage. We analysed the seasonal dynamics of the overall communities represented by these 3,090 vOTU genomes throughout the time-series. The Bray-Curtis dissimilarity index of all the possible pairs of samples was estimated and the average plotted as a function of the temporal distance between them (Fig. 2a). The significant sinusoidal pattern revealed peaks of dissimilarity at the 6-, 18-, and 32-month time gaps, while communities exhibit a maximum similarity at 0-, 12-, and 24-month time gaps between them. This remarkably stable annual fluctuation suggests that viral communities are strongly influenced by bottom-up processes, such as seasonality and the sensitivity of the host community to environmental fluctuations in the surface water.

**Figure 2.**
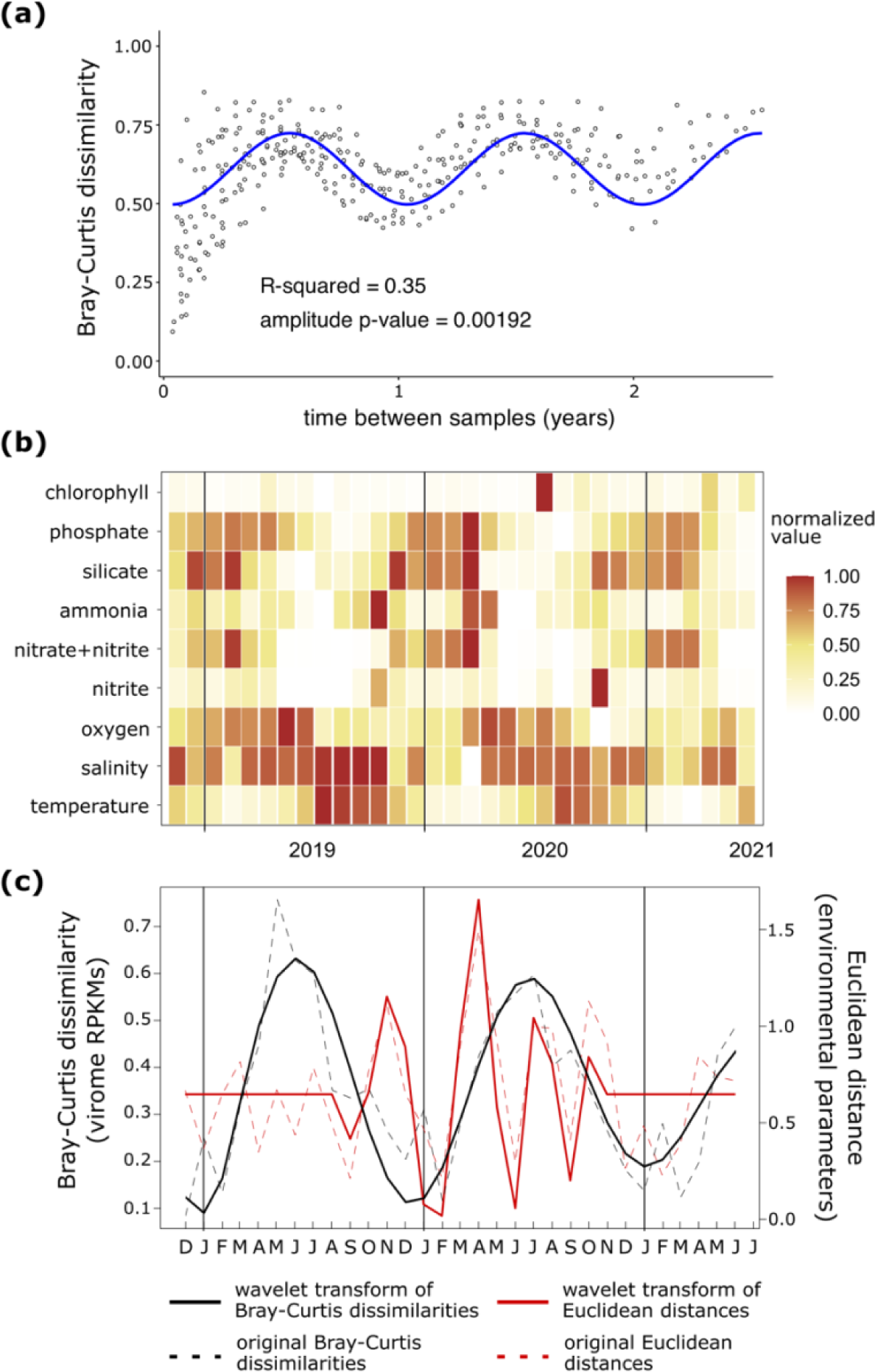
Bray–Curtis dissimilarity time decay analysis of the virome composition in the surface Western English Channel and its relationship with environmental changes. Pairwise Bray-Curtis dissimilarity in the viral community was estimated using the Reads Per Kilobase per Million mapped (RPKMs) of 3,090 representative genomes over a period of more than two and a half years. **(a)** Bray– Curtis dissimilarity time decay analysis. Bray–Curtis dissimilarities were averaged to establish a correlation with time distance (time gap between samples). A harmonic linear regression model was employed to identify significant seasonal trends in both the complete dataset and the analysed fractions. A harmonic linear regression model was employed to identify the significant seasonal trend (p < 0.05). **(b)** Normalized values (0–1) of the nine environmental parameters used to estimate the Euclidean distance between monthly consecutive samples. **(c)** Wavelet transforms (solid red line) of the monthly consecutive Euclidean distances representing the environmental changes and Bray–Curtis dissimilarities of the viral communities (solid black line). The original Euclidean distances and Bray-Curtis dissimilarities are shown as dashed lines, red and black respectively.

To interrogate whether this strong seasonal signal was driven by a few abundant genomes, we fractionated the high-quality populations based on the distribution of the calculated RPKMs. The first quartile, the interquartile interval and the third quartile were extracted, and Bray-Curtis dissimilarities were recalculated. The three fractions exhibited similarly remarkable annual stability, suggesting that seasonality is not dependent on abundance. Using the Fisher’s G test for time-series, 2,484 genomes were categorized as seasonal, and 606 as non-seasonal (Fig. S1). A sinusoidal harmonic linear regression was fitted to each of these categories to confirm that the signal was significantly correlated with the seasonal fraction Bray-Curtis dissimilarities. The Bray-Curtis dissimilarities between the seasonal genomes displayed a significant correlation to the sinusoidal regression, which was absent in the non-seasonal fraction (Fig. S1). Similarly, each of these groups were divided into the three quantiles described before. In the seasonal fraction, the seasonality signal was present in all quantile ranges evaluated, while absent in each of the non-seasonal quantiles. These temporal patterns were supported when the extended 26,851 vOTUs dataset was analysed (Fig. S2). These results suggest that viral communities are mostly composed of prevalent seasonal populations with a wide distribution of abundances, and a smaller fraction displaying irregular non-seasonal temporal patterns.

We evaluated how monthly environmental changes influence the rate of change of the virome composition. Consecutive Euclidean distances were generated based on nine environmental measurements (Fig. 2b). Wavelet transforms of the monthly consecutive Euclidean distances and Bray-Curtis virome composition dissimilarity were estimated and compared. Bray-Curtis virome dissimilarity showed an oscillatory trend in which the rate of change increases sharply from December-January to June-July. Meanwhile, this rate decreases from June-July to December-January (Fig. 2c). Euclidean distances did not show a similar oscillatory pattern. Wavelet coherence analysis confirmed that Euclidean and Bray–Curtis distances were out of phase during the studied period (Fig. S3).

### Viral populations that show similar patterns of longitudinal abundance can be clustered into highly defined chronotypes

Among the seasonal and non-seasonal groups of viral populations, different genomes exhibit comparable temporal patterns. To have a highly defined characterization of these patterns, we developed and employed a novel unsupervised machine learning method to *de novo* cluster them into chronotypes. Therefore, a chronotype is defined as a cluster of viral population representatives that have similar longitudinal abundance patterns and are optimally maximized to be different enough from other temporal clusters within the community.

Out of the 2,484 seasonal genomes, 108 optimal chronotypes were determined using this method. From the 606 non-seasonal genomes, 46 chronotypes were the optimal number (Fig. 3). Based on vConTACT2 taxonomic characterization, we observed that viral genera diversity increased linearly as a function of the number of members a chronotype has without reaching a plateau (Fig. S4). Both seasonal and non-seasonal chronotypes tend to be diverse following a linear relationship between the number of different taxonomic genus and number of members.

**Figure 3.**
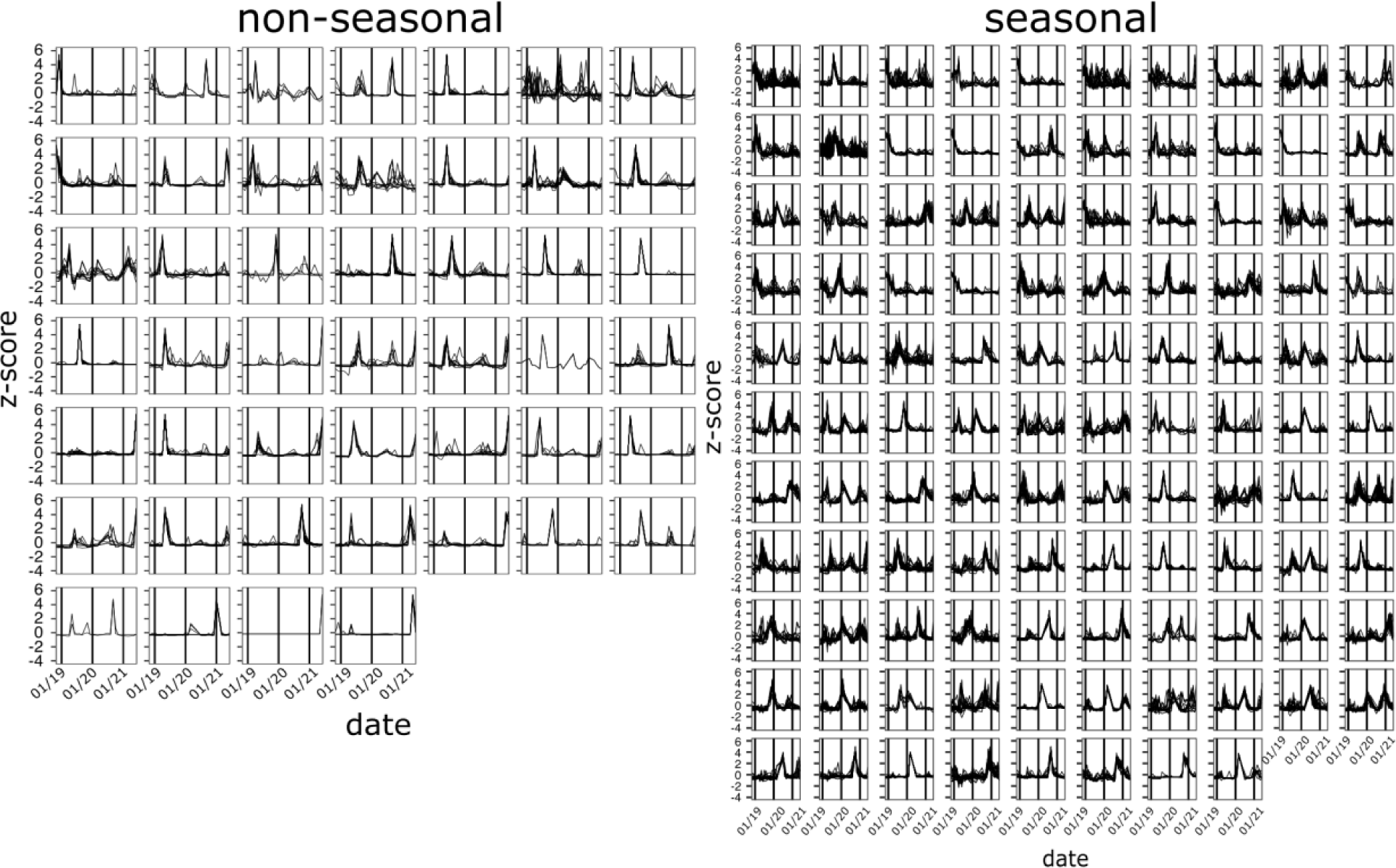
Seasonal and non-seasonal chronotypes derived from an unsupervised machine learning clustering method. Z-score normalized values of each of the 3,090 longitudinal RPKMS organized and plotted by its chronotype membership, represented by each panel. On the left side, 606 non-seasonal normalized RPKMS time-series are represented in 46 different chronotypes. On the right side, 2,484 seasonal normalized RPKMS time-series are represented in 108 chronotypes.

### Seasonal chronotypes show a significant enrichment in a subset of auxiliary metabolic genes associated with seasonal phages

We hypothesised that characteristic AMGs from vOTUs genomes associated with strongly seasonal hosts at the WEC (i.e *Synechococcus*) [38], such as cyanophages photosystem ll genes, would be enriched in seasonal chronotypes [79]. Out of the total 379 PHROG functional categories detected in all vOTU genomes, six showed a significant differential abundance between those in seasonal versus non-seasonal chronotypes: 2-OG(FeII) oxygenase superfamily, ferrochelatase, primase, nucleotide-sugar epimerase, amidoligase, and phosphoheptose isomerase (Fig. 4). These predicted proteins may be classified as AMGs, except primases which are categorized in the DNA, RNA, and nucleotide metabolism. 2-OG(FeII) oxygenase and ferrochelatase genes have been found to co-occur in pelagiphages genomes [66] and catalyze the oxidation of multiple compounds [80]. Nucleotide-sugar epimerase and phosphoheptose isomerase have been reported in cyanophages [81,82]. Amidoligases have been suggested to be involved in preventing superinfection by other phages in phage phiEco32 [83]. Contrary to our hypothesis, AMGs enriched in seasonal chronotypes did not include those associated with cyanophage photosystem manipulation.

**Figure 4.**
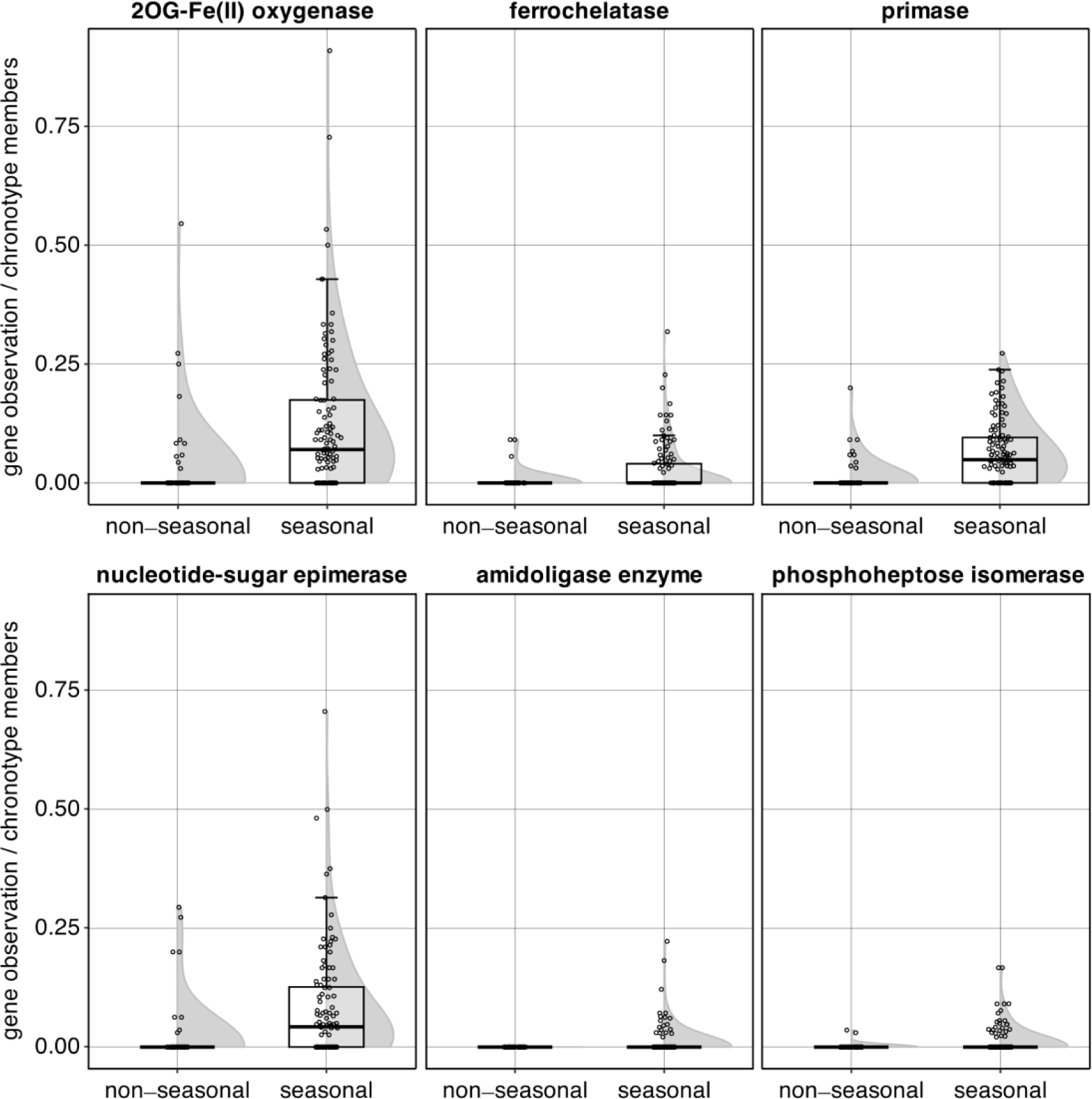
Seasonal chronotypes have a significant higher number of Auxiliary Metabolic Genes (AMGs). Distribution of normalized gene observations in seasonal and non-seasonal chronotypes of the significantly different functional categories. Violin plots and box plots are overlapped to show the distribution of the values. Gene categories are organized from left to right: (top) 2-OG(FeII) oxygenase (phrog61), ferrochelatase (phrog2954) and primase (phrog21866). (bottom) nucleotide-sugar epimerase (phrog540), amidoligase (phrog2443), and phosphoheptose isomerase (phrog1238).

To further investigate the relationship between the seasonality of enriched AMGs, their taxonomy, and phylogenetic relationships, we performed a maximum likelihood phylogenetic analysis of all the translated proteins of the coding sequences belonging to 2-OG(FeII) oxygenase superfamily, ferrochelatase, and primase categories (Fig. 5). These functional categories have representative genes in cultivated pelagiphage genomes, which allow us to accurately evaluate whether these genes occur broadly through the viral diversity or are a constrained genomic feature of abundant and seasonal pelagiphages. The 2-OG(FeII) oxygenase phylogenetic tree shows multiple lineages that might be related to the functional diversification of this family (Fig. 5a). Interestingly, 2-OG(FeII) oxygenases from cultured pelagiphages are distributed throughout many of these clades. Additionally, the metagenomic proteins derived from vOTUs that clustered with Pelagiphages (based on vConTACT2 results) are also broadly distributed through the phylogenetic tree. Similarly, ferrochelatases are mostly retrieved from vOTUs clustered with Pelagiphages and are broadly distributed in the phylogenetic tree (Fig. 5b). Primases are more conserved genes, and clades are not mostly derived from pelagiphage taxonomically related sequences (Fig. 5c). Importantly, the gene-sharing profile classification of metagenomic retrieved vOTUs with pelagiphages by vConTACT2 is not intended to extrapolate host prediction but provide taxonomical context of the genomes containing these sequences. Therefore, we can interpret that beyond the host, the expanded seasonal taxonomical groups related to the analysed pelagiphages are enriched with the three described gene categories.

**Figure 5.**
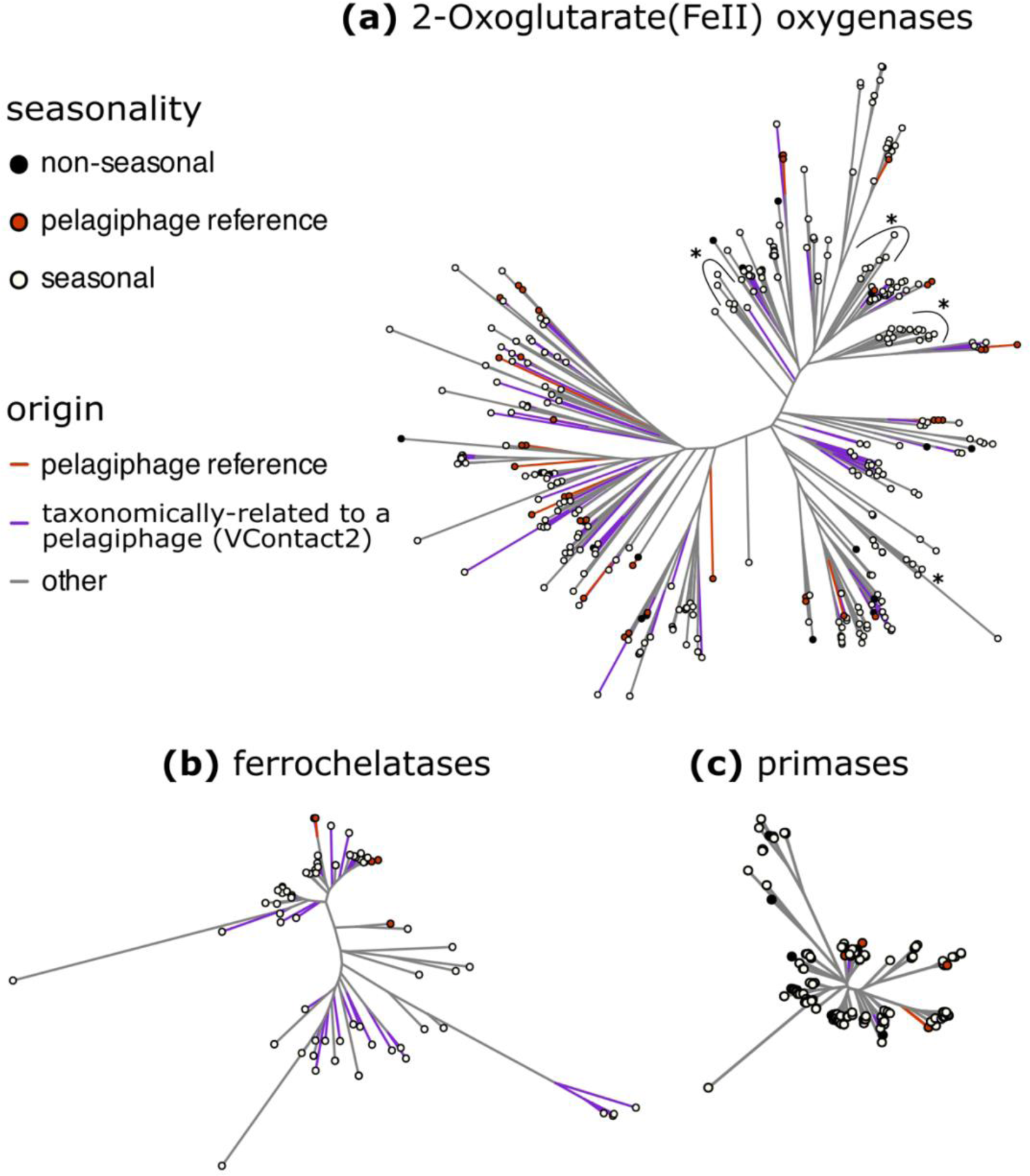
Phylogenetic analysis of the seasonal enriched gene categories. Maximum likelihood phylogenetic trees of the translated coding sequences of the genes driving the differences between seasonal and non-seasonal chronotypes. For all the phylogenetic trees, branches are colour-coded based on the vOTU where the predicted proteins were originated: red for the pelagiphage references, purple for vOTUs clustered with pelagiphage genomes (taxonomically-related to pelagiphages at the genus level) and gray for other (non-pelagiphage taxonomically related vConTACT2 cluster). The terminal nodes are overlapped with a colour-coded circle indicating the temporal pattern, seasonal vOTU or non-seasonal vOTU, of the genome which the coding sequence was retrieved or if it was retrieved from known pelagiphage genomes. Asterisks indicate clades without any taxonomically related pelagiphage origin protein. **(a)** 2OG-Fe(II) oxygenase **(b)** primase, and **(c)** ferrochelatase.

Specific chronotypes were enriched in genes involved in nucleotide metabolism, lysis, and structure (tail, head, and packaging proteins) (Fig. S5). Given their roles in essential viral functions, these enrichments, represented by the high-end outliers of the distributions, may just be coincidences of chronotypes having members with better characterized genes in the databases. The enrichment of seasonal chronotypes with 2-OG(FeII) oxygenases and co-occurring ferrochelatases (Fig. S6) underscores the viral adaptations and their influence on the seasonal cycle of their hosts by potentially expanding and manipulating their metabolic capacity. Furthermore, no specific AMGs were found that could broadly explain the differences between seasonal and non-seasonal vOTUs. Instead, enriched AMGs appeared to be associated with highly represented seasonal phages.

### Chronotypes are influenced by temperature, salinity, and nutrient concentrations

To evaluate the potential effects and influence of environmental conditions on the temporal patterns of seasonal and non-seasonal chronotypes, we identified the medoids for each chronotype and generate correlation profiles with the available surface water physico-chemical variables (Fig. 6). Based on the correlation profiles, two major groups were identified and separated by differential correlations with temperature salinity and nutrients. These two large trends are represented by clade b and c. Clade b chronotypes are positively correlated with temperature and salinity, while negatively with nutrients (NO_2_+NO_3_, PO_4_, SiO_2_). Clade c chronotypes are negatively correlated with temperature and salinity and positively with nutrients (NO_2_+NO_3_, PO_4_, SiO_2_). The presence of 2-OG(FeII) oxygenases and co-occurring ferrochelatases AMGs did not show an enrichment pattern in any of the previously described clades but were consistently found in seasonal chronotypes regardless of their relationship with environmental variables. While the identified chronotype groups exhibit varied correlations with temperature, dissolved oxygen, salinity and nutrients, the absence of a consistent correlation of the seasonal-enriched AMGs with the evaluated parameters suggests that the functional diversity of these genes allow them to be selected within multiple environmental conditions.

**Figure 6.**
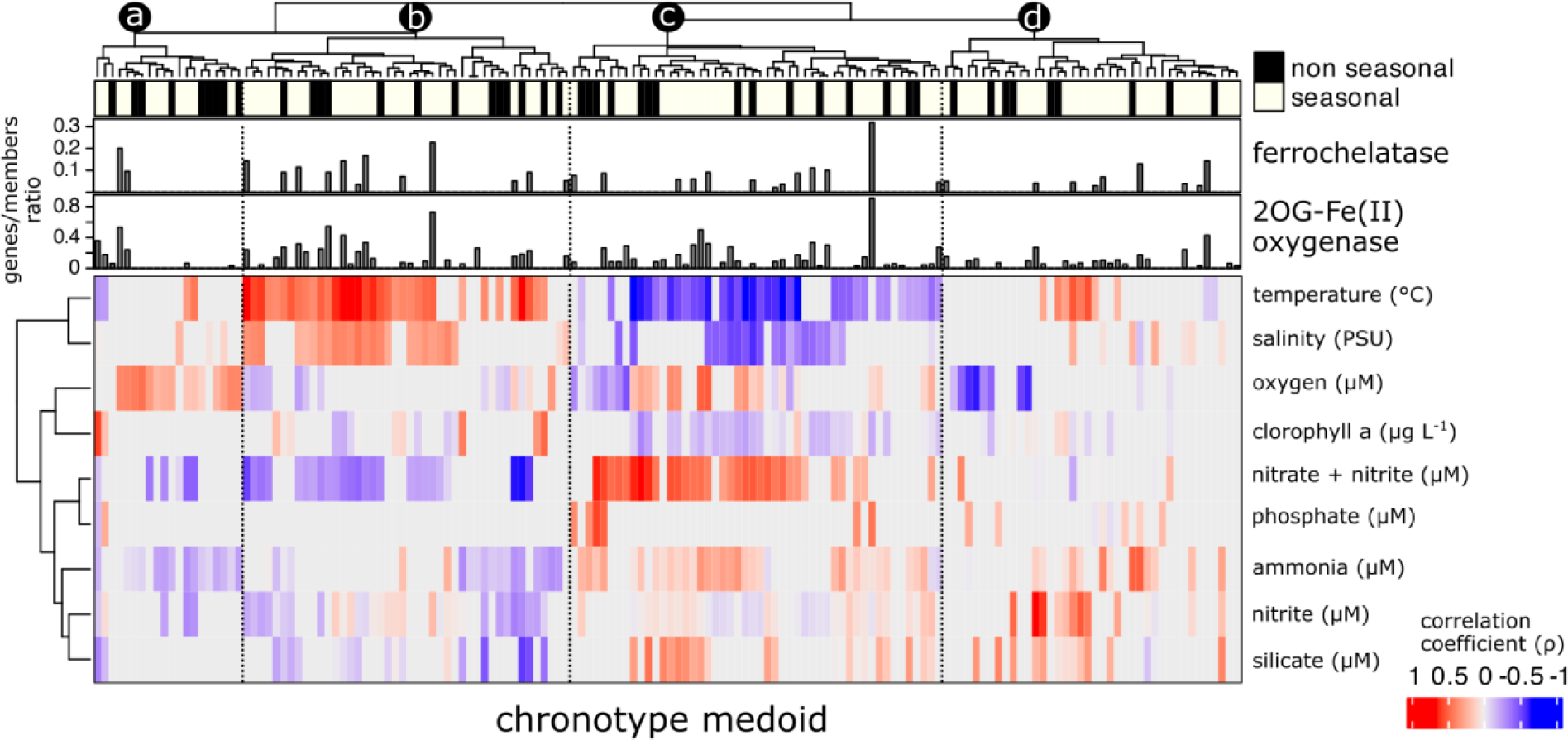
Environmental correlation analysis with medoid representatives of the chronotypes. The correlation heatmaps are color-coded if the correlation was significative (p<0.05). Correlation heatmap between chronotype medoids and oceanographic environmental parameters. The top dendrogram represent the medoid clustering based on the correlation profiles of the heatmap. The terminal node attached bar plot indicates the temporal pattern, seasonal or non-seasonal, of the medoid. Below, two bar plot panels depict the distribution of the auxiliary metabolic genes, ferrochelatase and 2OG-Fe(II) oxygenase, in the chronotype represented by the medoid. Based on the correlation profile, four groups of chronotypes are distinguished. “a” Medoids mostly correlate positive with oxygen concentration. “b” Medoids positively correlate with temperature and salinity, while negatively with nutrients (NO_2_+NO_3_, PO_4_, SiO_2_). “c” Medoids mostly correlate positively with nutrients (NO_2_+NO_3_, PO_4_, SiO_2_, NH_4_), while negatively correlated with temperature and salinity. “d” Medoids do not show a consistent pattern of correlations.

### Viral genomes undergo rapid polymorphism turnover, with a few seasonal chronotype members displaying annual recurrence of genetic variants

We sought to test whether the Red Queen dynamics shown in the relatively stable system such as SPOTS [27] would be evident in the WEC, which undergoes greater fluctuations in mixing and temperature over both short and long timescales [63]. Intra-population genotypic changes throughout the time-series were analysed by comparing the genetic polymorphic profiles of each vOTU. First, we analysed the polymorphic profiles of those vOTUs (>10kb) with a minimum coverage of 10X across 90% of its length throughout all the samples. This dataset was composed of 56 high quality vOTUs (4,472 SNPs). Pairwise comparison of population profiles at each month compared with previous and subsequent months showed that the genetic similarity, represented by the percentage of shared polymorphisms, decreases over time (Figs. 7,S7).

**Figure 7.**
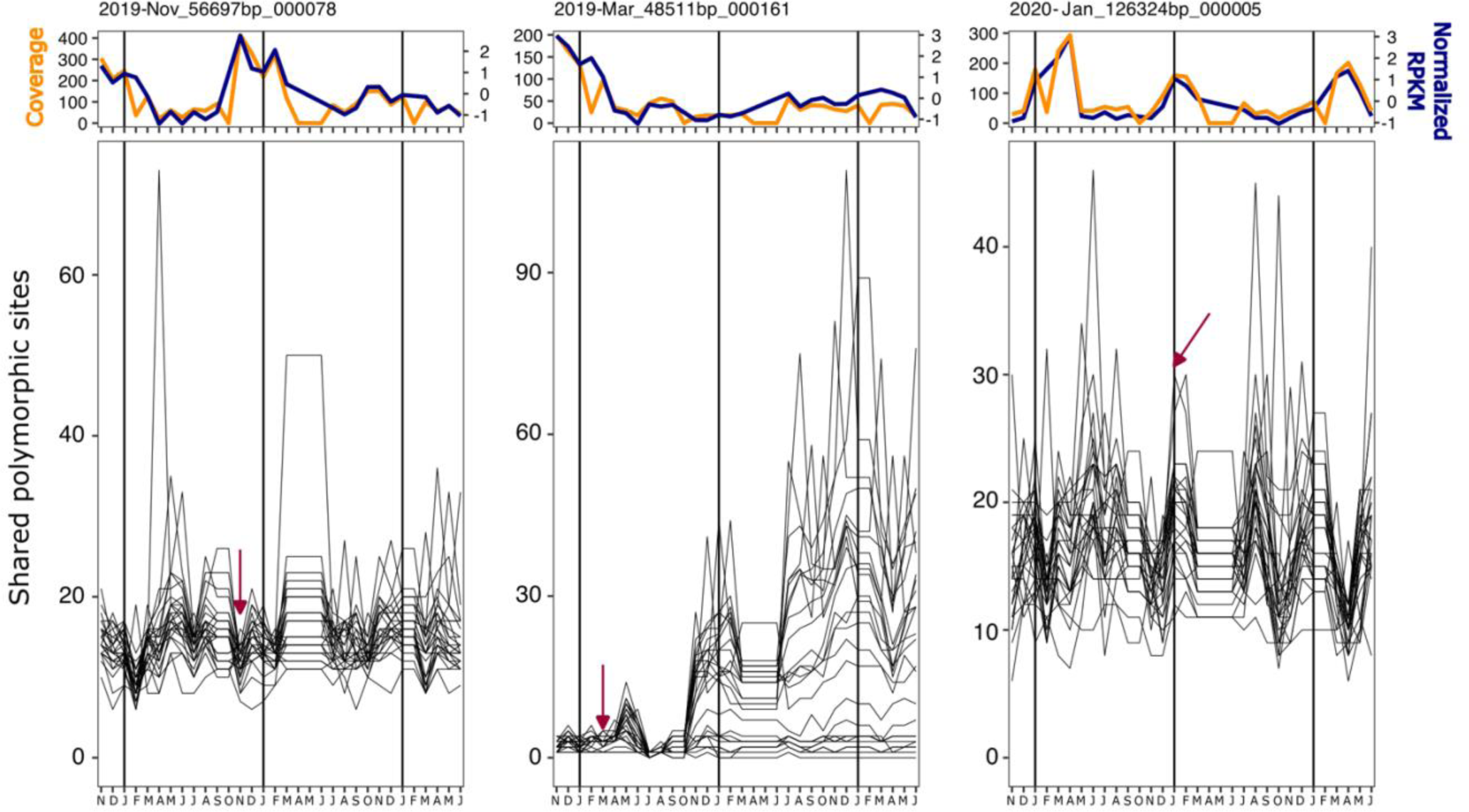
Temporal dynamics of polymorphic profiles of ubiquitous and abundant vOTUs. Three representative examples out of the 56 vOTUs that fulfilled the coverage, breadth, and identity requirements in the analysis are shown. Each vOTU belongs to a different chronotype, which abundance pattern is shown on the top panels by (1) coverage (in yellow and fixed in the left y-axis) and (2) z-score normalized RPKM (in blue and fixed in the right y-axis). A pairwise comparison of shared polymorphic sites is shown in the bottom panels. Each line represents the collection of comparisons of a fixed month to the rest of the profiles. A red arrow indicates the sample of origin of the representative vOTU sequence.

Even though variance density correlates positively with coverage and RPKM (Fig. S8), when considering high-quality vOTUs with a minimum coverage of 10X across 90% in at least one sample, we recovered variation profiles from most of the vOTUs comprising the total of our curated dataset (2,997 out of 3,090; ∼97%). From these, we selected the vOTU members (n=365) of 16 chronotypes that exhibited the clearest seasonality (similar abundances recurrence), either annually peaking on summer or winter. As expected, the variant density of these vOTUs correlated significantly with their RPKM and coverage (Figs. S9,S10,S11), but it allowed us to explore whether genetically similar subpopulations may reoccur in a similar trend as their chronotype seasonality. By estimating the mean of the percentage of shared polymorphisms of a specific vOTU in all the potential time difference comparisons (0 to 32 months) we tested which vOTUs polymorphic profiles display a seasonal signal (Figs. 8,S12,S13). We determined that the subpopulation genetic profiles of 31.5% vOTUs of the seasonal chronotypes show a significant partial recurrence of genetic profiles that peaks in the 12 months pairwise comparisons. However, this signal fades out in profile comparisons of 16 months or more. We compared this pattern to vOTUs belonging to sporadic non-seasonal chronotypes. These are characterized by presenting a sharp peak of abundance at a particular month, without any similar recurrence within the time-series. We selected the vOTU members (n=261) of 16 sporadic non-seasonal chronotypes and performed the same analysis. Only 16% of these vOTUs showed a seasonal signal, and 12-month recurrence median was less than half of that in the recurrent seasonal profiles. This low recurrence of genotype profiles of a few sporadic vOTUs represent a residual or background signal which might not be filtered in the Fisher G-test of seasonality. These results support that a large fraction of vOTUs is subjected to a constant turnover of genetic population variants consistent with Red Queen Dynamics. However, a smaller fraction of vOTUs display patterns in which profiles of polymorphic populations are similar when time-periods of 12 months are compared.

**Figure 8.**
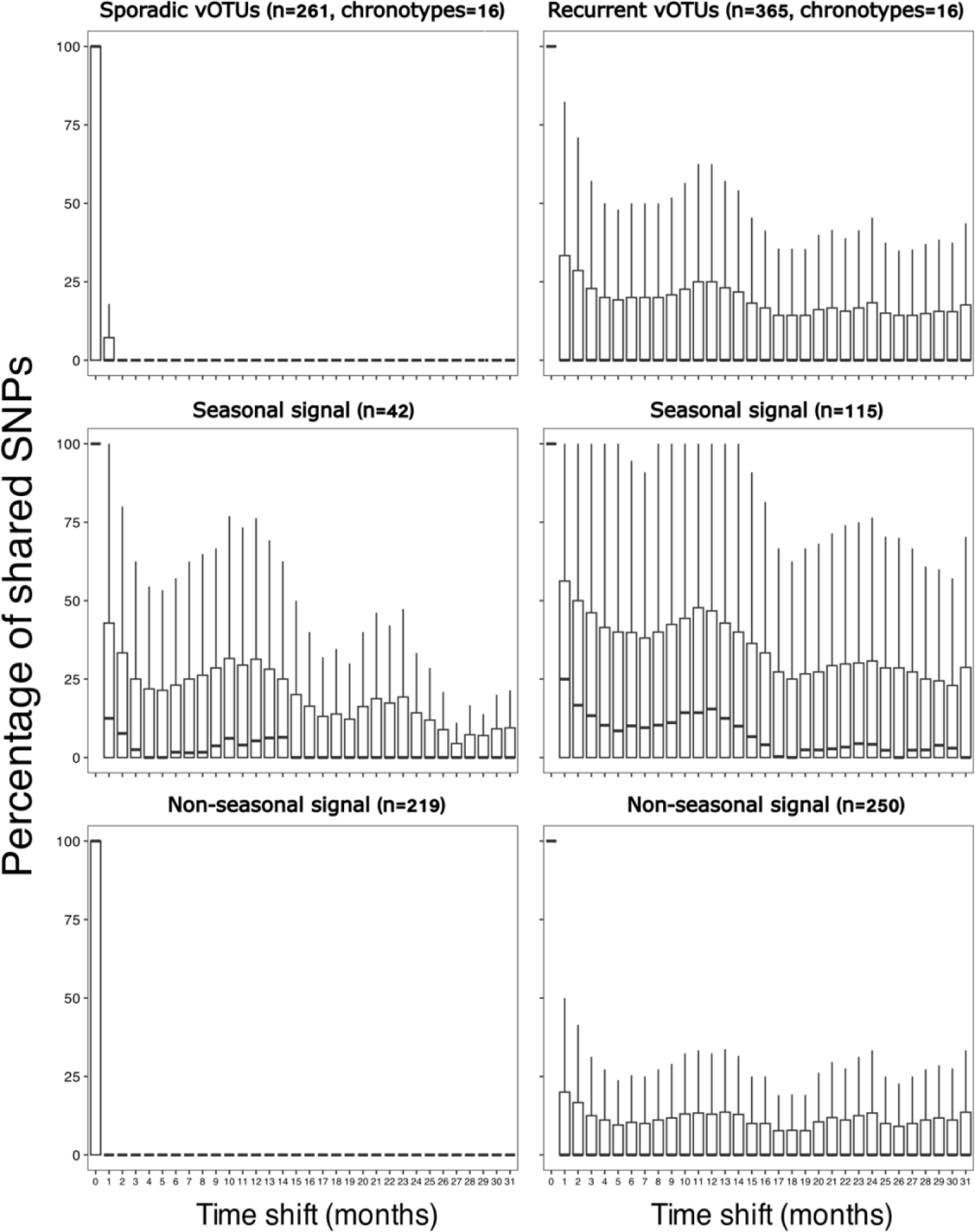
Polymorphic profile comparisons in non-seasonal sporadic and seasonal annual recurrent vOTUs. 16 non-seasonal (left panels) and 16 seasonal chronotypes (right panels) were selected and the polymorphic profiles of its member vOTUs were compared pairwise between all potential combinations of the 27 available samples. The percentage mean of each unique monthly comparisons (from 0 to 32 months) of shared SNPs for each vOTU was estimated. The resulting distribution of means were then plotted as whisker plots (top panels). Using Fisher’s G test for time-series, we determined which vOTUs have a distribution of shared means with a seasonal signal (mid panels). The vOTUs without a significant seasonal signal (bottom) are plotted for comparison purposes. In all the panels, outliers of the distributions were omitted to highlight the differences in the majority of the data.

## Discussion

In this study, we investigated the genomic commonalities and differences of viral populations that covaried through time at the WEC and the environmental factors shaping them. Our sampling strategy focused on the “free” viral size fraction (<0.2μm) at the surface (0-5m), which is assumed to predominantly recover lytic dsDNA viruses [28]. This was confirmed in this study. Overall, the viral community composition showed a remarkably stable annual periodicity at the WEC. A similar pattern was first reported at the San Pedro Ocean Time-Series [27], and later observed in Osaka Bay [29] where it was described that this annual trend was significantly correlated with the prokaryotic community composition and the seasonal environmental variables. A cycle of high-rate compositional changes from winter to summer and steadier transitions of viromes from summer to winter was described at WEC. As this cycle did not follow the same trends in environmental fluctuations, we hypothesize these oscillations are mostly driven by changes in host composition and productivity.

To conduct a detailed analysis of the temporal patterns followed by vOTU genomes, we developed an unsupervised machine learning clustering method to classify the viral populations into chronotypes based on the comparisons of the distances between the individual abundance time-series. This agnostic approach overcomes the major challenge of determining a meaningful threshold for partitioning the covariances by integrating an iterative search of an optimal number of clusters (*k*). Our method provides important benefits - the most important the automatic determination of optimal clusters based on distances that allow us to capture local and global patterns. From the user perspective, multiple advantages exist such as: interpretability, scalability, flexibility in data types, and few pre-processing requirements. Unsupervised machine learning distance-based clustering of time series provides a powerful method for classifying high-throughput datasets without prior knowledge of the classes [84].

The chronotype framework has proven successful in delineating the diversity of co-occurring populations, identifying genes involved in seasonal dynamics, and assessing the influence of environmental conditions on the assembly of these populations. Chronotypes did not exhibit specific taxonomic enrichments, possibly indicating a linear relationship with the diverse number of hosts that follow similar seasonal patterns of abundance. Further analysis, integrating host-derived data (such as amplicon or cellular fraction metagenome-assembled genomes) with viral chronotypes, or an improvement in accuracy of host-prediction from phage genomes [85], may offer better insights into these seasonal relationships and mechanisms of interaction.

AMGs play a crucial role in expanding host metabolic capacity and enhancing virus production in marine ecosystems [86,87]. These genes are fundamental genomic signatures across various locations and depths [28,88–91] (Hurwitz et al., 2015; Hurwitz et al., 2016; Coutinho et al., 2017; Luo et al., 2020; Coutinho et al., 2023). Here, we demonstrate that a subset of AMGs is enriched in seasonal chronotypes. Notably, 2-OG(FeII) oxygenases and ferrochelatases often co-occur in pelagiphages from the Podoviridae family [66]. Phylogenetic analysis suggests that most of these predicted proteins have an origin in pelagiphage taxonomically related vOTUs. SAR11 ecotypes show consistent seasonal patterns in 16S rRNA ASVs at the WEC [63], which might explain an important fraction of the viral gene enrichments found in this study. The presence of these genes across seasonal chronotypes, with different patterns of correlation with environmental parameters, suggests a conserved enrichment in pelagiphages and taxonomically related vOTUs regardless of the season at which the specific host thrives. 2-OG(FeII) oxygenases catalyse diverse biochemical reactions, contributing to metabolite diversity and participate in sulphur and phosphate metabolism in microbial systems [80,92–94]. Because of their vast phylogenetic diversity, the specific functions and their role in seasonal virus-host interactions remain to be determined.

It has been previously shown that ocean viral populations showed a rapid and constant turnover of their genetic variant profiles as a result to overcome the mutual selective pressure that viruses and their hosts exert on each other, suggestive of Red Queen dynamics [27]. Here we analysed whether this pattern was consistent in the WEC. Our results showed that most of the viral populations followed a similar trend, in which a specific polymorphism profile increases to a maximum and then decays within a few months. However, when we subset and analyse the most consistent seasonal chronotypes (annual recurrent with similar abundances than previous years), we found that some populations have a strong recurrent signal of a shared collection of polymorphisms when profiles 12-months apart are compared. This signal fades out as comparisons are done for longer periods of time. These results suggest that some phage populations might have a slower range of change, specifically in an annual scale instead of monthly, although still consistent with Red Queen dynamics as these similarities faded out as the years progressed.

This study presents an innovative framework to interrogate the temporal patterns of viruses in the surface ocean. Our findings reveal that the stable seasonality observed in the viral community can be deconstructed into seasonal chronotypes and a smaller fraction of non-seasonal patterns. We suggest that the seasonal patterns of viral chronotypes are adapted to couple their productivity with that of their recurrent seasonal hosts, such as SAR11 and *Synechococcus*, while sporadic chronotypes experience an increase in abundance triggered by specific conditions, leading to short-term ‘boom and burst’ events, likely involving copiotrophic hosts. Both mechanisms broadly follow the Kill-the-Winner model [95], but operate at different time scales. The significant association of some AMGs with seasonal chronotypes suggests that a diverse array of versatile functions within these genes have been evolutionarily selected as an adaptive strategy to sustain the replication and production of viruses at a seasonal scale. The robustness of the time-series clustering method introduced in this study will facilitate the generation of highly defined co-occurring longitudinal units in other molecular datasets. As demonstrated by this study, such an approach will contribute to uncovering novel ecological and evolutionary patterns in longitudinal data.

## Acknowledgements

The authors thank the crew of the Plymouth Marine Laboratory vessel ‘Quest’ for collection of seawater samples. Research was supported by high performance computing resources provided by the University of Exeter.

## Author Contributions

**Luis M. Bolaños:** conceived and designed the experiments, analysed the data, prepared figures and/or tables, authored and reviewed drafts of the paper, and approved the final draft.

**Michelle Michelsen:** performed the experiments, authored, and reviewed drafts of the paper, and approved the final draft.

**Ben Temperton:** conceived and designed the experiments, analysed the data, prepared figures and/or tables, authored and reviewed drafts of the paper, and approved the final draft.

## Data Availability

Short read viromes are deposited in NCBI under the Bioproject PRJNA804019.

