## Supplemental Material for "Viral chronotypes and their role in shaping seasonal viral dynamics in the Western English Channel"

### Supplementary Material

#### Supplementary methods:

**Unsupervised machine learning framework to generate chronotypes: ChronoClustR.**
We present an unsupervised machine learning framework for generating chronotypes, named ChronoClustR. Time-series clustering is an essential framework across multiple disciplines and has undergone constant development (Aghabozorgi *et al.*, 2015). In the era of big data, unsupervised machine learning clustering methods have played a fundamental role in generating patterns and classes when no a priori knowledge is available. In this study, we propose an alternative method to generate highly defined clusters of time-series datasets (Fig. 1). The following pipeline was developed in R v4.3.

**User input:**
This pipeline requires an equidistant set of time-series with no missing values. The user should input the following:

1. A data frame of the vOTU RPKM (or any other value) by timepoint (sample).
2. Metadata with the corresponding date.
3. Number of bootstrap iterations (this value should be greater than 1).
4. Size of the subsample (default = 100).
5. The starting date.
6. The end date.
7. Specify the frequency or number of timepoints.

**Processing of individual replicates:**
**Subsampling:** Randomly subsample (without replacement) the collection of time-series using the user-specified size. The user may consider estimating a subsample size that generates an exact integer of subsamples (to cover all time-series within the dataset) or the one that generates the smallest residue (Fig. 1, step 6).

**Unsupervised Clustering of Each Subsample** (Default Size 100): A function named "centroid_generator" clusters each subsample using the TSclust algorithm (Fig. 1, step 7) with the following default parameters:

1. Type (clustering method): Partitional
2. k (number of clusters to evaluate): 3 to (n-1)
3. Distance: Euclidean
4. Centroid: Partitioning around medoids (PAM)

Any of the above parameters can be adjusted to fulfil user requirements. The options are those used by the TSclust algorithm (Montero P. and Vilar J.A., 2015). After generating the clustering of each subsample, the "centroid_generator" function will retrieve the best number of clusters based on the maximum Silhouette score for each tested number of clusters (*k*). This function will then recalculate the clusters for this *k* and retrieve a list of centroids and their associated cluster members.

**Unsupervised Clustering of Centroids and Optimized *k*:** The centroids generated from the clusters of all subsamples are extracted and concatenated. This collection of centroids is clustered using the k-means algorithm, testing all potential numbers of clusters (*k*) from 1 to n-1. The generated within-cluster sum of square for each computed cluster for each value of *k* is extracted (Fig. 1, step 8). Based on the curve generated from the relationship between the total within-cluster sum of squares and *k*, we define the optimal number of clusters by estimating the closest value of *k* to the first derivative (slope of the tangent line). This first derivative is calculated on the fitted curve estimated on the relationship between the number of clusters and its within-cluster sums of squares (Fig. 1, step 9).

**Unsupervised clustering with an optimized *k*:** Once the best *k* is estimated based on the centroids of the subsamples, this *k* is used to generate three types of clustering (k-means, hierarchical clustering, and TSclust) on the original dataset (Fig. 1, step 10).

**Bootstrap:** The above steps are coded to be repeated as many times as indicated by the user (Fig. 1, step 11). Warning: As the number of iterations increase, the memory needed to generate the co-occurrence matrix also increases, limiting the successful analysis by the memory capacity of the user's computing infrastructure.

**Co-occurrence matrix and chronotype generation**: After *n* replicates of bootstrap, the configurations of the different clusters of each replicate are used to generate a co-occurrence matrix (Fig. 1, step 12). The user can choose to use any of the three clustering methods from step 10. The co-occurrence values are normalized to the number of iterations (*n*) used for the bootstrap. Finally, a dendrogram is generated using the hclust function, and the maximum optimal number of clusters from all the iterations is used to partition the dendrogram. Each partition will be considered a chronotype (Fig. 1, step 13).

R scripts and data used to generate chronotypes are available at https://github.com/lbolanos32/WEC_Chronotypes_2024.

#### Supplementary Figures and Tables:

##### Table S1: Environmental metadata and virome accession (raw short reads).

| Sample | Date | Date_wav | SRA (NCBI) | Temp (°C) | Fluor (V) | Dens (kg.m3) | Sal (PSU) | O2 (uM) |
|---|---|---|---|---|---|---|---|---|
| 2018-Nov | 20/11/2018 | 07/11/2018 | SRR18164412 | 13.3915 | 0.7567 | 1026.437 | 35.1443 | 245.262 |
| 2018-Dec | 04/12/2018 | 07/12/2018 | SRR18164411 | 12.1169 | 0.7576 | 1026.2895 | 34.6282 | 253.638 |
| 2019-Jan | 14/01/2019 | 07/01/2019 | SRR18164400 | 11.011 | 0.6565 | 1026.7145 | 34.9094 | 260.321 |
| 2019-Feb | 11/02/2019 | 07/02/2019 | SRR18164392 | 9.3312 | 0.434 | 1026.4479 | 34.1984 | 270.148 |
| 2019-Mar | 11/03/2019 | 07/03/2019 | SRR18164391 | 10.0049 | 0.4015 | 1026.9742 | 35.0154 | 267.477 |
| 2019-Apr | 01/04/2019 | 07/04/2019 | SRR18164390 | 10.3946 | 0.4577 | 1026.947 | 35.067 | 270.54 |
| 2019-May | 07/05/2019 | 07/05/2019 | SRR18164389 | 11.4882 | 0.3845 | 1026.701 | 35.0051 | 289.791 |
| 2019-Jun | 03/06/2019 | 07/06/2019 | SRR18164388 | 13.6776 | 0.2984 | 1026.3111 | 35.0588 | 278.34 |
| 2019-Jul | 22/07/2019 | 07/07/2019 | SRR18164387 | 17.33 | 0.5402 | 1025.6144 | 35.2189 | 224.859 |
| 2019-Aug | 12/08/2019 | 07/08/2019 | SRR18164386 | 16.9253 | 0.4441 | 1025.7414 | 35.2582 | 242.563 |
| 2019-Sep | 02/09/2019 | 07/09/2019 | SRR18164410 | 16.3641 | 0.5957 | 1025.8784 | 35.2647 | 239.814 |
| 2019-Oct | 02/10/2019 | 07/10/2019 | - | 16.1277 | 0.6609 | 1025.9259 | 35.2569 | 229.279 |
| 2019-Nov | 05/11/2019 | 07/11/2019 | SRR18164409 | 13.7882 | 0.6881 | 1025.7409 | 34.3507 | 228.511 |
| 2019-Dec | 02/12/2019 | 07/12/2019 | SRR18164408 | 12.0838 | 0.4746 | 1026.4812 | 34.8668 | 237.124 |
| 2020-Jan | 20/01/2020 | 07/01/2020 | SRR18164407 | 9.8135 | - | 1026.43 | 34.2763 | 239.545 |
| 2020-Feb | 05/02/2020 | 07/02/2020 | SRR18164406 | 9.7321 | - | 1026.5673 | 34.4347 | 240.232 |
| 2020-Mar | 02/03/2020 | 07/03/2020 | SRR18164405 | 9.1438 | - | 1026.0072 | 33.5967 | 265.805 |
| 2020-Apr | 07/04/2020 | 07/04/2020 | - | 9.9763 | - | 1026.9247 | 34.9476 | 283.251 |
| 2020-May | 04/05/2020 | 07/05/2020 | - | 11.1393 | 0.7241 | 1026.7412 | 34.9736 | 276.215 |
| 2020-Jun | 08/06/2020 | 07/06/2020 | - | 12.9412 | 0.9564 | 1026.4234 | 35.0108 | 256.979 |
| 2020-Jul | 07/07/2020 | 07/07/2020 | SRR18164404 | 13.731 | 3.657 | 1026.2237 | 34.9597 | 271.245 |
| 2020-Aug | 10/08/2020 | 07/08/2020 | SRR18164403 | 16.5421 | 1.6689 | 1025.6532 | 35.0254 | 255.713 |
| 2020-Sep | 07/09/2020 | 07/09/2020 | SRR18164402 | 15.9492 | 0.9631 | 1025.7824 | 35.0157 | 235.425 |
| 2020-Oct | 12/10/2020 | 07/10/2020 | SRR18164401 | 14.7043 | 1.6795 | 1025.8094 | 34.6917 | 201.806 |
| 2020-Nov | 09/11/2020 | 07/11/2020 | SRR18164399 | 13.6133 | 1.5318 | 1026.2712 | 34.9894 | 214.929 |
| 2020-Dec | 07/12/2020 | 07/12/2020 | SRR18164398 | 11.9147 | 0.6191 | 1026.5668 | 34.9362 | 220.044 |
| 2021-Jan | 11/01/2021 | 07/01/2021 | SRR18164397 | 9.5617 | 1.0781 | 1026.7136 | 34.5838 | 236.779 |
| 2021-Feb | 08/02/2021 | 07/02/2021 | - | 9.0447 | 0.7727 | 1026.5613 | 34.2846 | 234.979 |
| 2021-Mar | 08/03/2021 | 07/03/2021 | SRR18164396 | 8.4951 | 0.7257 | 1026.8666 | 34.5622 | 226.851 |
| 2021-Apr | 13/04/2021 | 07/04/2021 | SRR18164395 | 9.334 | 1.0157 | 1027.0553 | 34.9701 | 248.859 |
| 2021-May | 12/05/2021 | 07/05/2021 | SRR18164394 | 10.8083 | 0.9175 | 1026.8078 | 34.983 | 259.301 |
| 2021-Jun | 07/06/2021 | 07/06/2021 | SRR18164393 | 14.2321 | 1.6498 | 1025.5136 | 34.1774 | 233.462 |

**Table S1 (continuation)**

| Sample | $NO_2$ (uM) | $NO_3$+$NO_2$ (uM) | $NH_4$ (uM) | $SiO_4$ (uM) | $PO_4$ (uM) | Chl *a* (ugL$^{-1}$) |
|---|---|---|---|---|---|---|
| 2018-Nov | 0.15 | 3.51 | 0.19 | 2.66 | 0.4 | 0.75 |
| 2018-Dec | 0.26 | 7.5 | 0.35 | 4.66 | 0.45 | 0.61 |
| 2019-Jan | 0.13 | 7.53 | 0.19 | 4.02 | 0.5 | 0.29 |
| 2019-Feb | 0.3 | 11.8 | 0.3 | 4.82 | 0.57 | 0.47 |
| 2019-Mar | 0.36 | 6.88 | 0.1 | 2.67 | 0.52 | 0.43 |
| 2019-Apr | 0.28 | 5.75 | 0.26 | 1.9 | 0.55 | 1.75 |
| 2019-May | 0.0000001 | 0.11 | 0.24 | 0.51 | - | 0.78 |
| 2019-Jun | 0.0000001 | 0.1 | 0.04 | 0.15 | - | 0.77 |
| 2019-Jul | 0.0000001 | 0.03 | 0.0000001 | 1.3 | - | 0.08 |
| 2019-Aug | 0.0000001 | 0.0000001 | 0.28 | 1.03 | 0.03 | 0.61 |
| 2019-Sep | 0.11 | 0.13 | 0.36 | - | 0.06 | 0.38 |
| 2019-Oct | 0.68 | 1.49 | 0.66 | 2.43 | 0.24 | 0.42 |
| 2019-Nov | 0.32 | 8.03 | 0.34 | 4.82 | - | 0.37 |
| 2019-Dec | 0.15 | 6.54 | 0.08 | 3.49 | 0.53 | 0.24 |
| 2020-Jan | 0.16 | 8.95 | 0.09 | 4 | 0.53 | 0.46 |
| 2020-Feb | 0.14 | 9.87 | 0.22 | 3.87 | 0.51 | 0.58 |
| 2020-Mar | 0.24 | 12.32 | 0.59 | 5 | 0.71 | - |
| 2020-Apr | 0.21 | 3.05 | 0.54 | 0.55 | 0.41 | - |
| 2020-May | 0.0000001 | 0.0000001 | 0.0000001 | 0.45 | 0.19 | 0.24 |
| 2020-Jun | 0.03 | 0.14 | 0.0000001 | 0.56 | - | 0.33 |
| 2020-Jul | 0.17 | 0.64 | 0.14 | 0.55 | 0.05 | 7.39 |
| 2020-Aug | 0.0000001 | 0.0000001 | 0.13 | 1.43 | 0.0000001 | 0.77 |
| 2020-Sep | 0.13 | 0.89 | 0.27 | 1.88 | 0.11 | 2.02 |
| 2020-Oct | 1.04 | 4.02 | 0.33 | 4.2 | 0.26 | 1.21 |
| 2020-Nov | 0.17 | 5.08 | 0.14 | 3.93 | 0.37 | 0.73 |
| 2020-Dec | 0.1 | 5.93 | 0.09 | 3.19 | 0.4 | 0.67 |
| 2021-Jan | 0.12 | 9.37 | 0.06 | 3.67 | 0.48 | 0.97 |
| 2021-Feb | 0.22 | 9.97 | 0.07 | 4.03 | 0.54 | 0.56 |
| 2021-Mar | 0.29 | 9.93 | 0.28 | 3.34 | 0.52 | 0.85 |
| 2021-Apr | 0.27 | 4.45 | 0.18 | 1.41 | 0.36 | 4.06 |
| 2021-May | 0.02 | 0.05 | - | 0.74 | 0.1 | 0.83 |
| 2021-Jun | 0.02 | 0.04 | 0.12 | 0.35 | 0.03 | 2.6 |

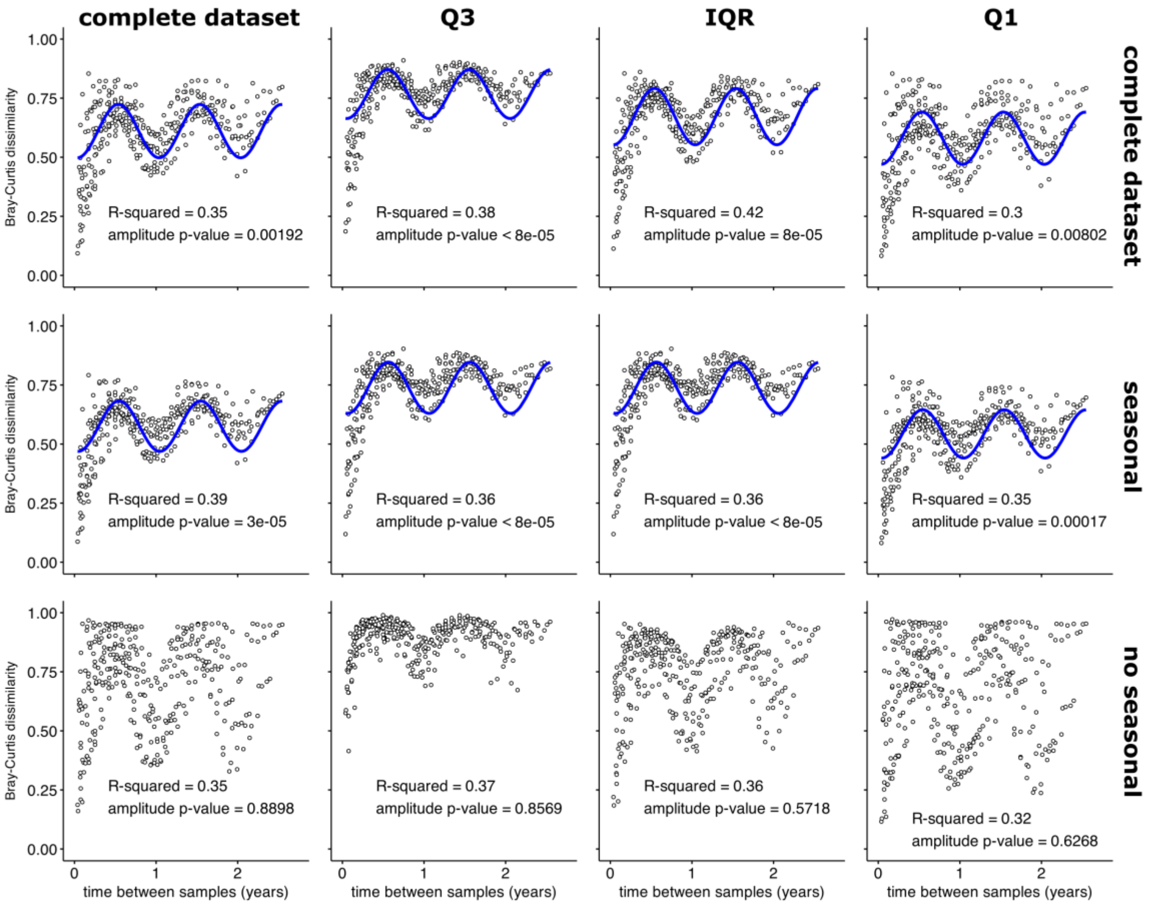


**Figure S1: Bray-Curtis dissimilarity time decay analysis of the high-quality virome composition in the Surface Western English Channel.** Pairwise dissimilarity in the viral community was estimated using the Reads Per Kilobase per Million mapped reads (RPKMs) of 3,090 representative genomes over a period of more than two and a half years. The Bray-Curtis dissimilarities were averaged to establish a correlation with time distance (time gap between samples). A harmonic linear regression model was employed to identify significant seasonal trends in both the complete dataset and the analysed fractions. If the Bray-Curtis dissimilarity sinusoidal trend was statistically significant ($p < 0.05$), the linear regression was illustrated in blue. Columns are organized from left to right based on abundance, with the first column representing the complete dataset, followed by the third quartile, the interquartile range, and the first quartile of the RPKM distribution of the 3,090 representative genomes. Rows are organized from top to bottom, depicting the complete dataset (top), the seasonal fraction (middle), and the non-seasonal fraction (bottom) of the RPKM distribution of the 3,090 representative genomes.

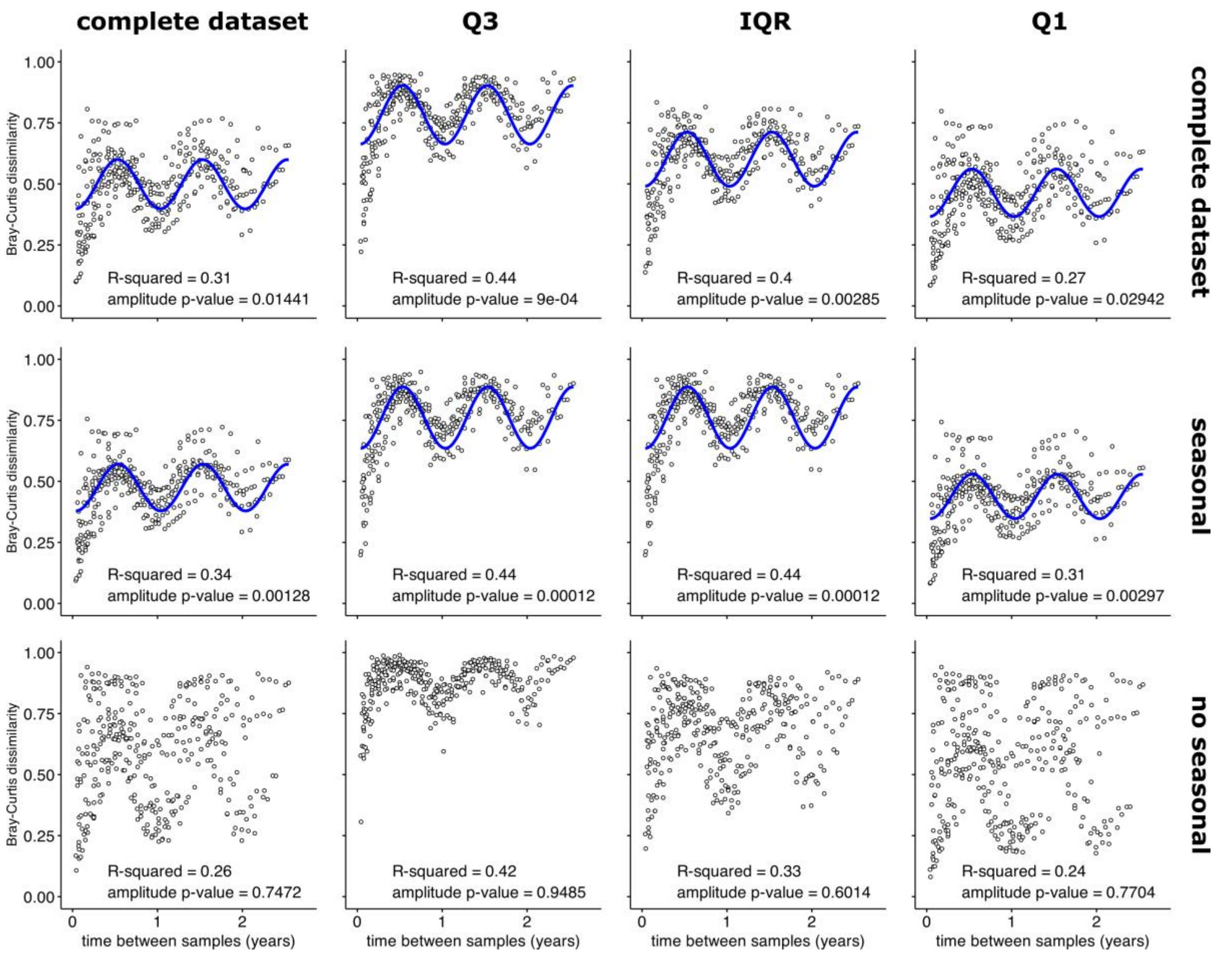


**Figure S2.- Bray-Curtis dissimilarity time decay analysis of the extended virome composition from the surface Western English Channel.** Pairwise viral community dissimilarity was estimated using the RPKMS of the 26,851 population representative genomes over more than two years and a half. Identically to Figure S2, Bray-Curtis dissimilarities were averaged to generate a one-to-one correspondence between dissimilarity and time distance (time gap between samples). A harmonic linear regression model was used to determine seasonal trends in the complete dataset and the analysed fractions. If the Bray-Curtis dissimilarity sinusoidal trend was significant ($p < 0.05$), the linear regression was plotted in blue. Columns are organized from left to right in abundance terms, being the first the complete dataset and followed by the third quartile, the interquartile range and the first quartile of the RPKM distribution of the extended population representative genomes. Rows are organized from top to bottom: the complete dataset (top), the seasonal fraction (middle), and the non-seasonal fraction (bottom) of the RPKM distribution of the extended population representative genomes.

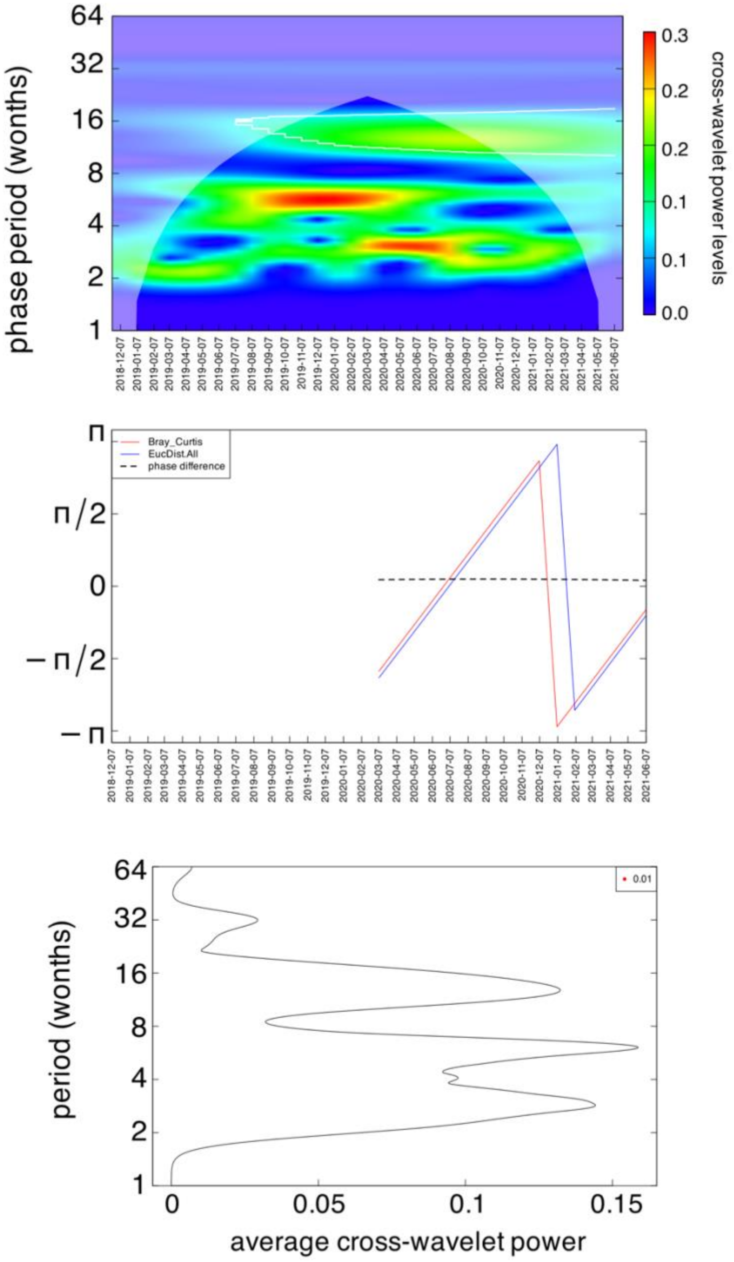


**Figure S3.- Wavelet coherence analysis between Bray-Curtis consecutive dissimilarities and Euclidean distances. (a)** Cross-wavelet power levels represented as a colour gradient. Periods with significant coherence ($p < 0.05$) are illustrated by a white line. **(b)** In phase representation of the two variables (red and blue) overlapped by the phase difference progression (dashed black line). **(c)** Average coherence for the significant in phase period ($p < 0.01$) is indicated in red.

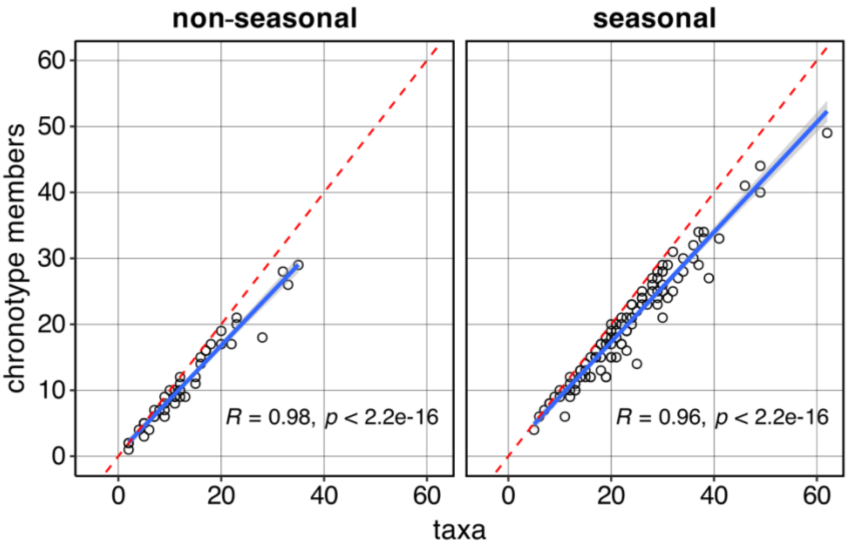


**Figure S4.- Viral Genus accumulation curves.** The taxonomic classification of the chronotype members were determined using gene-sharing profiles (vConTACT2 generated). Left panel shows the seasonal chronotypes and the right panel the non-seasonal. The dashed red line indicates a x=y linear relationship as reference. A significant positive linear correlation (spearman) exists between the number of vOTUs, and the number of genera represented in each chronotype.

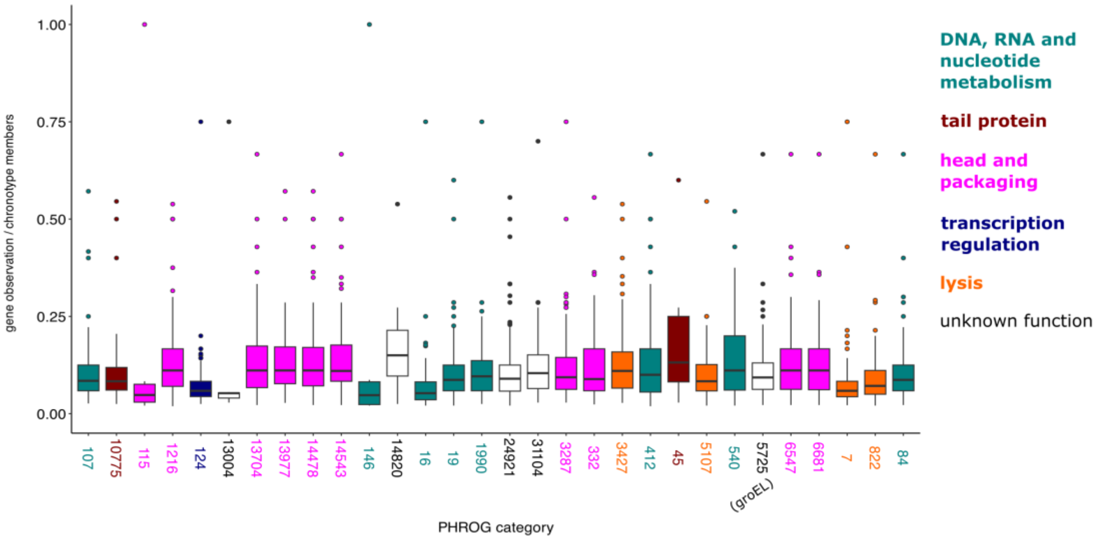


**Figure S5.- Proteins enriched in different chronotypes mostly represent structural functions and nucleic acids metabolism.** Distribution of normalized gene observations in all chronotypes organized by PHROG category. Only distributions of PHROG categories with at least a chronotype with a 0.5 ratio value of genes:chronotype members are shown. We set the enrichment threshold to be theoretically half of the chronotype members having a gene belonging to the specific displayed category. 30 PHROG categories fulfil this requirement. The distribution of these categories is color-coded based on the higher-rank PHROG functional characterization. All values over 0.5 represent outliers in their distribution, confirming this threshold as a meaningful.

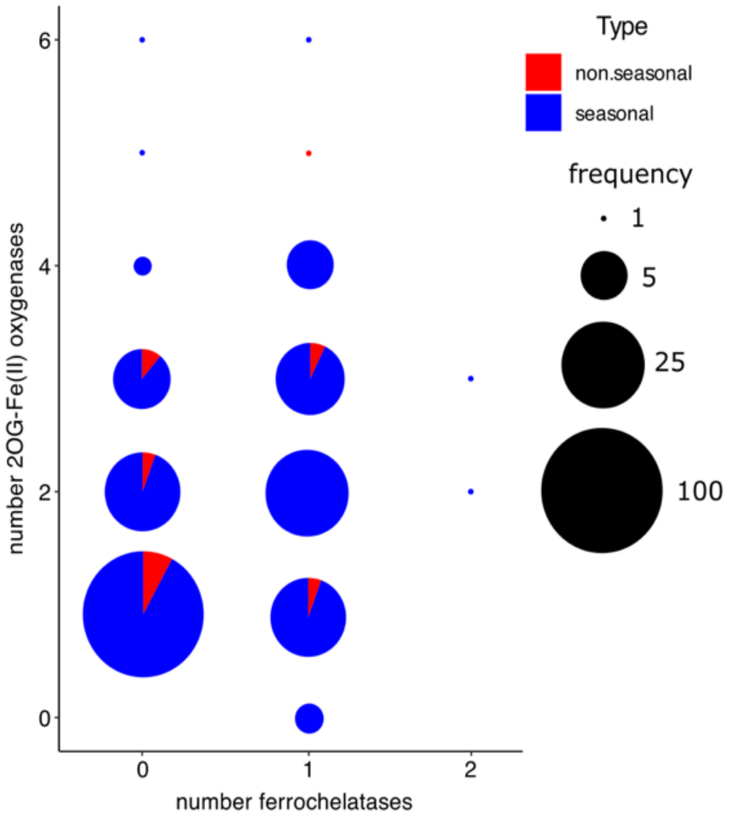


**Figure S6.- Co-occurrence of 2OG-Fe(II) oxygenase and ferrochelatase on the same population representative genome.** This plot represents the frequency of all pairwise combinations in the number of co-occurring 2OG-Fe(II) oxygenase and ferrochelatase coding sequences. Frequency is represented by the pie-chart size, while the percentage of seasonal and non-seasonal chronotypes that contribute to this frequency is represented by the color-coded fractions of the pie-charts.

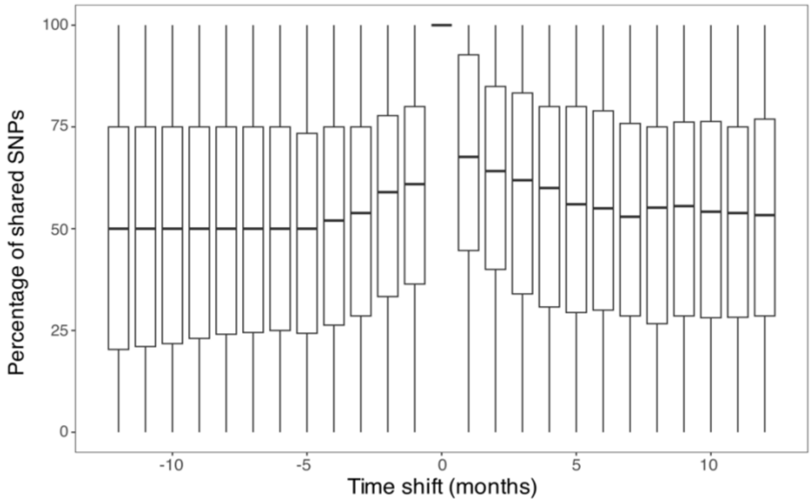


**Figure S7.- Comparison of the polymorphic profiles of the ubiquitous and abundant vOTUs vOTUs with a minimum coverage of 10X across 90% in all samples.**

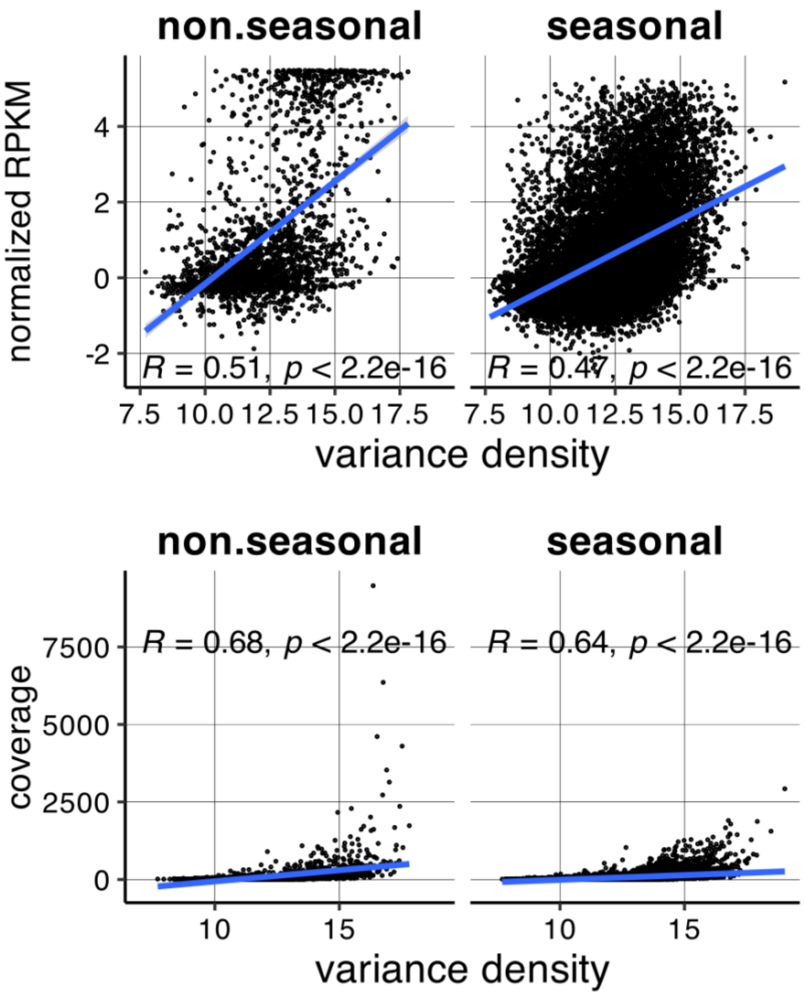


**Figure S8.- Correlation of variance density with normalized RPKMs and coverage of the high-quality vOTUs with a minimum coverage of 10X across 90% in at least one sample.**

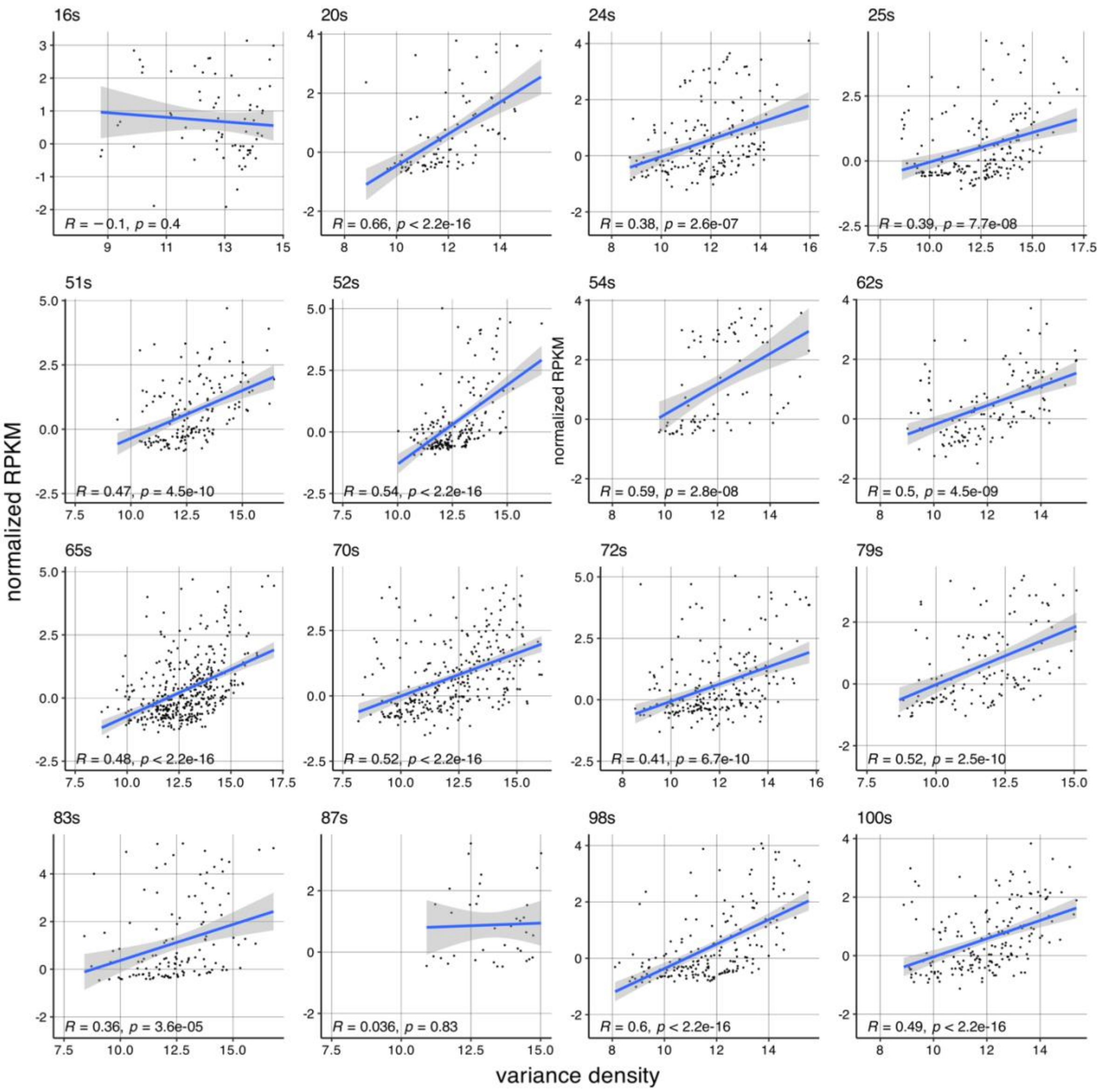


**Figure S9.- Correlation of variance density with normalized RPKMs of the vOTUs from the selected seasonal annual recurrent chronotypes.**

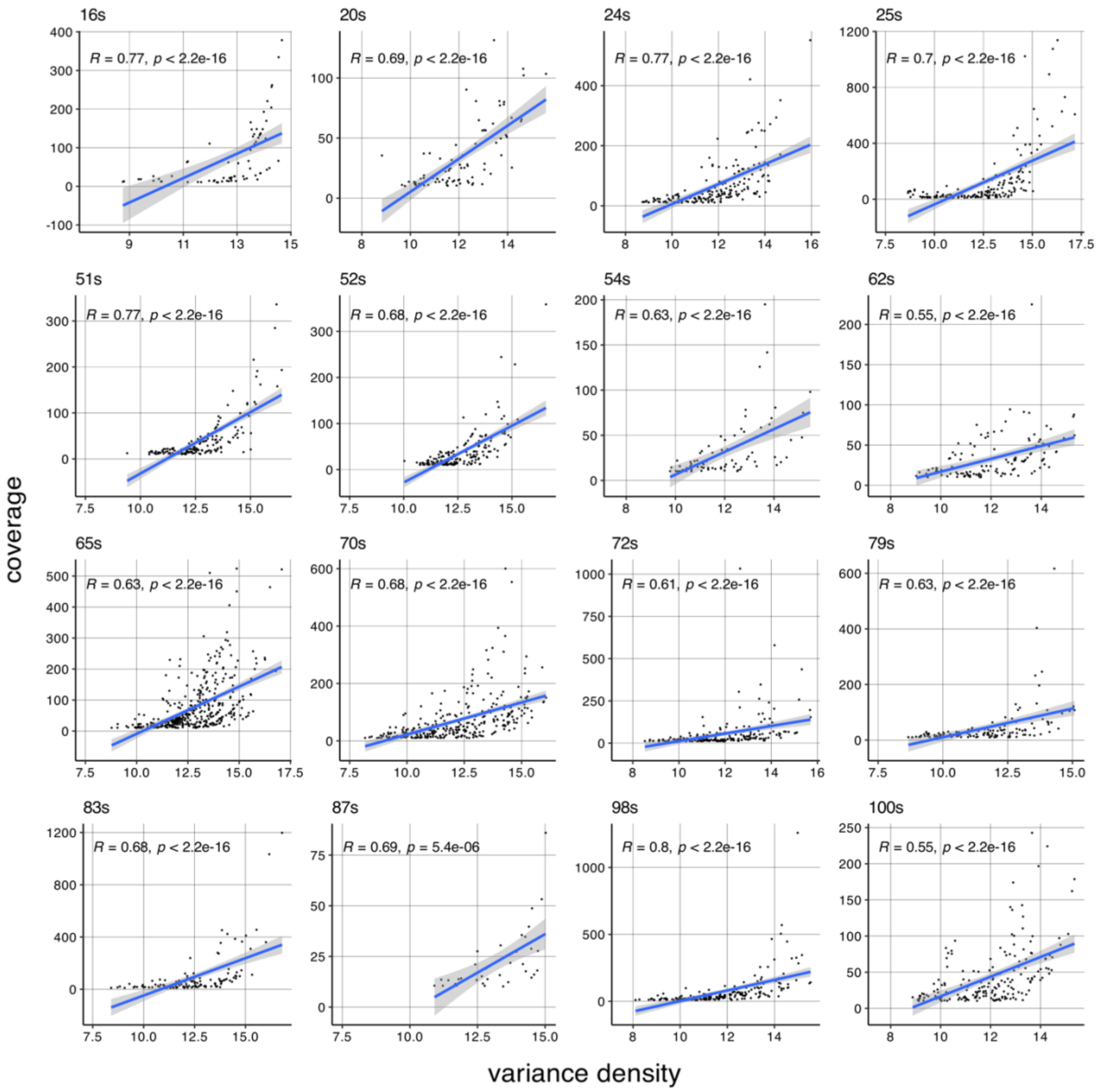


**Figure S10.- Correlation of variance density with coverage of the vOTUs from the selected seasonal annual recurrent chronotypes.**

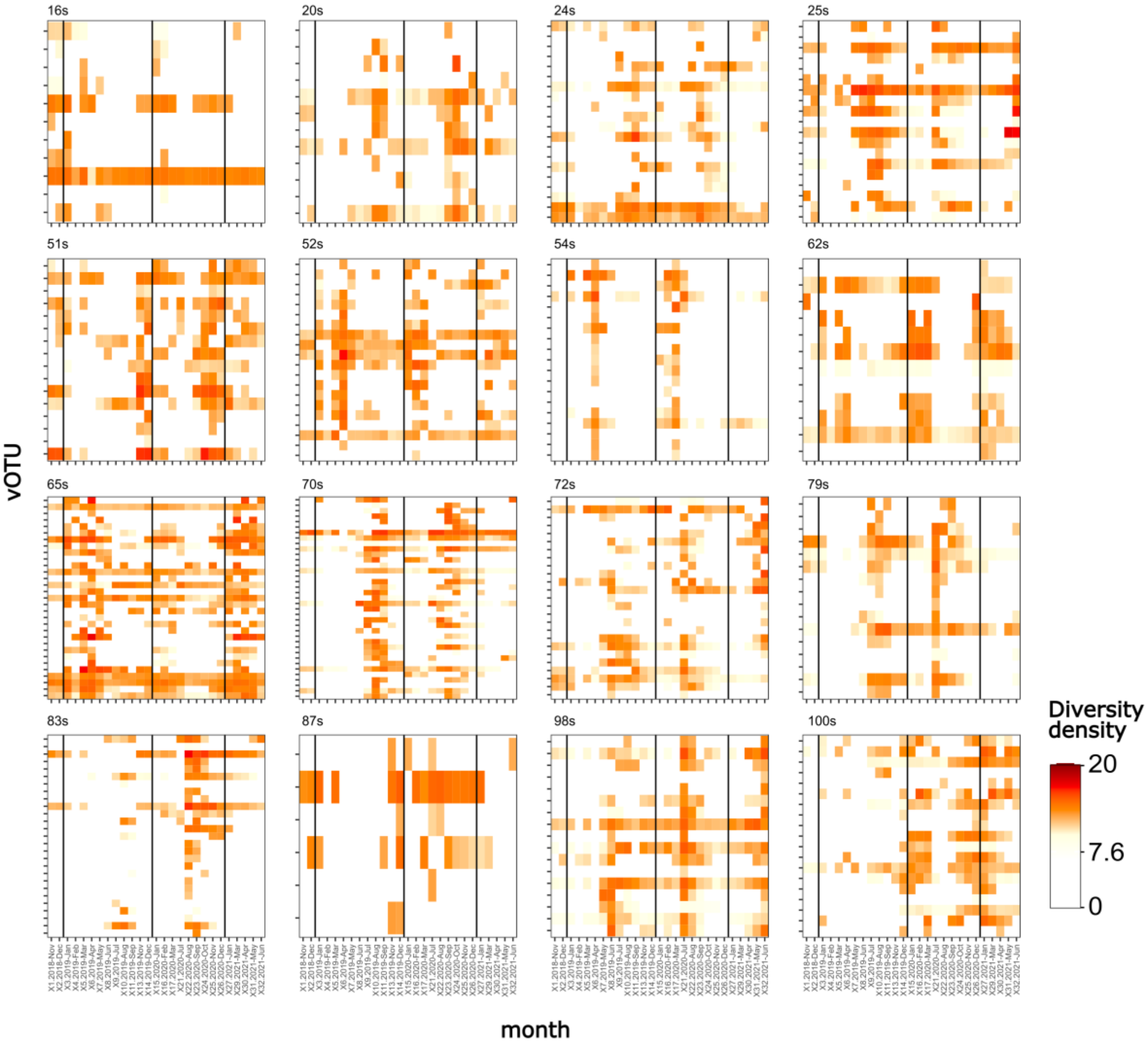


**Figure S11.- Variance density throughout the time series of the vOTUs from the selected seasonal annual recurrent chronotypes.** Each row represents a vOTU member of the chronotype (organized by panels). The diversity density is temporally sorted (x-axis).

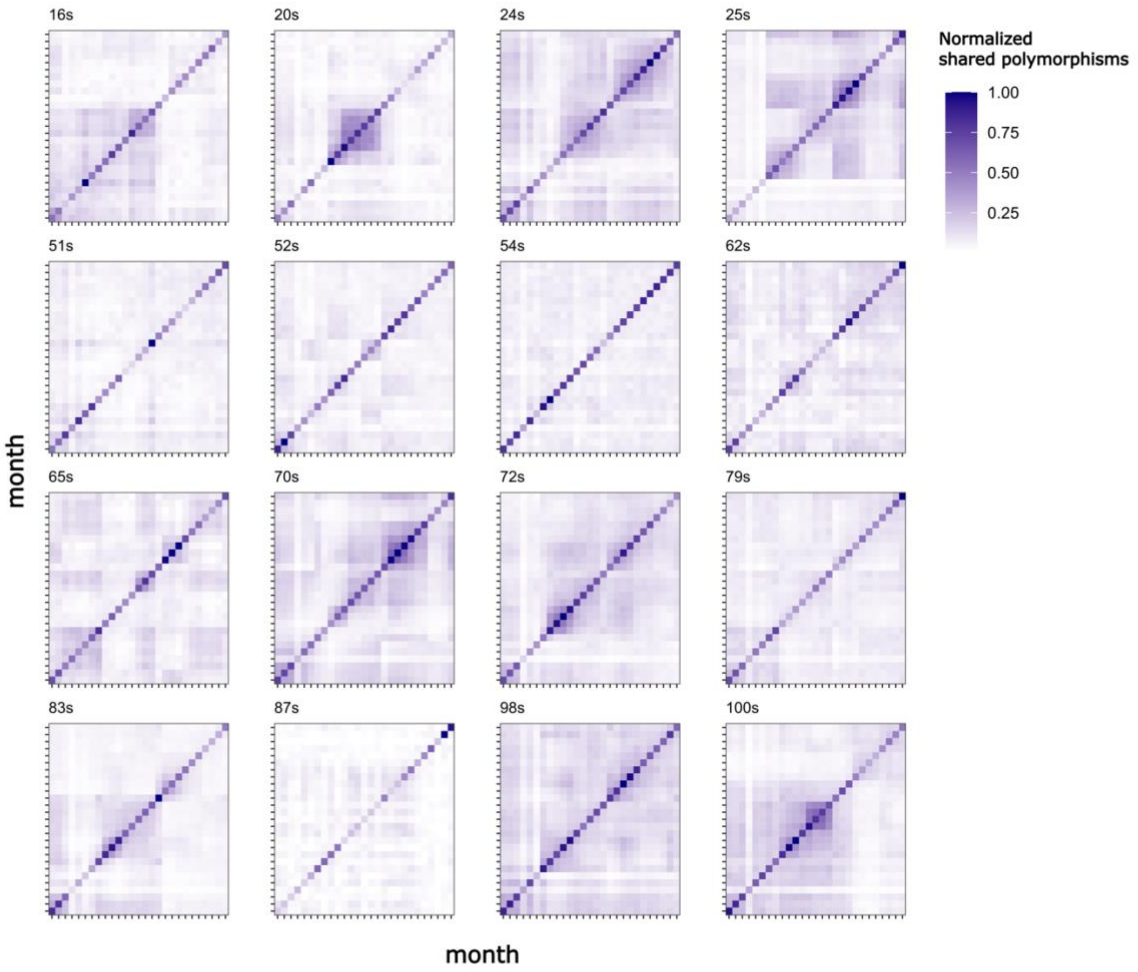


**Figure S12.- Monthly pairwise comparison of the normalized shared polymorphisms by chronotype.** For each chronotype we estimated the shared polymorphisms of all its member vOTUs in a pairwise comparison (all the potential combination of months). Each resulting chronotype matrix was normalized by setting the maximum value as 1 (feature scaling).

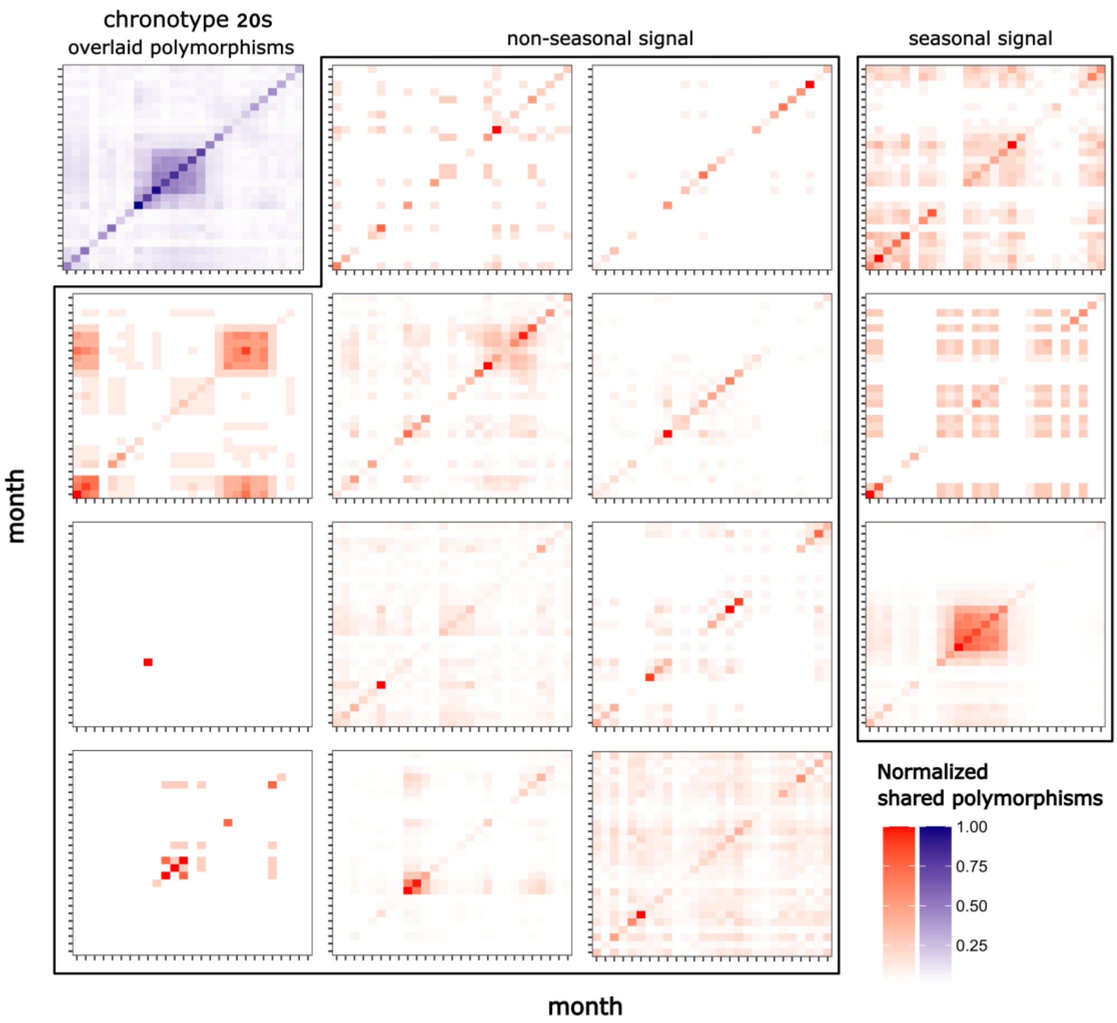


**Figure S13.- Chronotype 20s vOTUs with seasonal and non-seasonal recurrent polymorphic positions. This example shows how the vOTU members of a chronotype can have multiple patterns of shared polymorphisms.** As in Fig. S12, the resulting matrices were normalized by setting the maximum value as 1 (feature scaling). The top right corner represents the merged chronotype profile of shared polymorphisms (as shown in Fig. S12). The rest of the panels are the individual vOTU profiles from where the top right panel was derived. vOTU panels were organized by placing those with no-seasonal signal on the left and the three vOTUs with a seasonal signal on the right.
